# FUS post-transcriptional splicing is autoregulated via RNA condensation with therapeutic potential for ALS-FUS

**DOI:** 10.1101/2025.02.01.633781

**Authors:** Wan-Ping Huang, Vedanth Kumar, Karen Yap, Haiyan An, Sabin J. John, Rachel E. Hodgson, Anna Sanchez Avila, Emily Day, Brittany C.S. Ellis, Tek Hong Chung, Jenny Lord, Michaela Müller-McNicoll, Eugene V. Makeyev, Tatyana A. Shelkovnikova

## Abstract

Mutations in the *FUS* gene cause aggressive and often juvenile forms of amyotrophic lateral sclerosis (ALS-FUS). In addition to mRNA, the *FUS* gene gives rise to a partially processed RNA with retained introns 6 and 7. We demonstrate that these FUSint6&7-RNAs form nuclear condensates scaffolded by the highly structured intron 7 and associated with nuclear speckles. Using hybridization-proximity labelling proteomics, we show that the FUSint6&7-RNA condensates are enriched in splicing factors and the m6A reader YTHDC1. These ribonucleoprotein structures facilitate post-transcriptional FUS splicing and depend on m6A/YTHDC1 for their maintenance. FUSint6&7-RNAs become hypermethylated in cells expressing mutant FUS, leading to their enhanced condensation and consequently, splicing. We further demonstrate that FUS protein is repelled by m6A. Thus, ALS-FUS mutations may cause an abnormal activation of FUS post-transcriptional splicing via altered RNA methylation. Strikingly, ectopic expression of FUS intron 6&7 sequences dissolves the endogenous FUSint6&7-RNA condensates, downregulating FUS mRNA and protein. Overall, we describe an RNA condensation-dependent mechanism regulating FUS splicing that can be harnessed for developing new therapies.

## Introduction

FUS is an abundant RNA-binding protein (RBP) with numerous roles in cellular RNA metabolism (Ratti and Buratti, 2016). The *FUS* gene located on chromosome 16 encodes a 526 amino acid protein. Since the discovery of FUS’s association with amyotrophic lateral sclerosis (ALS) in 2009 (Kwiatkowski et al., 2009; Vance et al., 2009), >50 ALS-FUS mutations have been identified, most of which are missense mutations (reviewed in Moens et al., 2025). Although FUS mutations account for only 4% of familial ALS (fALS), they are the most frequent cause of juvenile ALS – a particularly aggressive form of the disease (Baumer et al., 2010; Deng et al., 2014; Grassano et al., 2022). The majority of FUS mutations map to its C-terminus and lead to the impairment or complete loss of the nuclear localization signal (NLS) (Moens et al., 2025). FUS cytoplasmic mislocalisation likely triggers a combination of loss- and gain-of-function mechanisms, however their relative contribution to ALS pathology is still debated. Studies in mice have demonstrated that mutant FUS expression, but not FUS knockout, is sufficient to cause neurodegeneration (Devoy et al., 2017; Korobeynikov et al., 2022; Lopez-Erauskin et al., 2018; Scekic-Zahirovic et al., 2016; Sharma et al., 2016), suggesting that FUS toxic gain of function plays a key role in the disease. This mechanism is further supported by the effects of a recently developed antisense oligonucleotide (ASO) therapy, where simultaneous depletion of both mutant and normal FUS inhibited neurodegeneration in mice and potentially in humans (Korobeynikov et al., 2022).

FUS accumulation in large cytoplasmic inclusions (FUS proteinopathy) in ALS-FUS postmortem tissue (Hewitt et al., 2010; Lashley et al., 2011; Nolan et al., 2016), ALS mutations in FUS 3’UTR leading to its overexpression (Sabatelli et al., 2013), and FUS mRNA upregulation in physiological cell models of ALS-FUS (An et al., 2019) all point to a disrupted control over its cellular levels as a disease hallmark. One reported mechanism supports changes in autoregulation, where FUS protein suppresses normal splicing of its pre-mRNA by promoting exon 7 skipping and nonsense mediated decay (NMD) (Zhou et al., 2013). A more recent study showed that introns 6 and 7 are often retained in FUS pre-mRNA (Humphrey et al., 2020). These highly conserved introns have multiple FUS protein binding sites, raising a possibility that FUS promotes their retention, thereby reducing the translatable FUS mRNA pool via a negative-feedback mechanism (Humphrey et al., 2020). These introns were found to be more efficiently spliced in *in vitro* and *in vivo* models of ALS-FUS (Humphrey et al., 2020). Intriguingly, FUS intron 6 and 7 retention was enhanced in a mouse ALS-FUS model upon introduction of the full human *FUS* transgene and was associated with a dramatic rescue of the disease phenotype (Sanjuan-Ruiz et al., 2021).

Given that modulation of disrupted FUS autoregulation may constitute an attractive therapeutic strategy, bypassing the undesirable FUS loss-of-function effects of the ASO treatment, we sought to refine our understanding of the molecular mechanisms underlying this process. We find that FUS transcripts with retained introns 6 and 7 (FUSint6&7-RNA thereafter) form a novel type of ribonucleoprotein condensate in the nucleus. Compositional analysis of these structures by a hybridisation-proximity labelling approach revealed enrichment of proteins involved in RNA splicing and regulation by N6-methyladenosine (m6A) modification. Mechanistic studies have led us to propose a role for this m6A-maintained condensate in FUS post-transcriptional splicing and a gain-of-function mechanism of mutant FUS in its dysregulation. Strikingly, introduction of exogenous FUS intron 6 and 7 sequences disrupted the condensation of endogenous FUSint6&7-RNA and depleted (mutant) FUS mRNA and protein. Our study advances current understanding of FUS expression regulation and describes a homeostatic approach to the modulation of FUS levels.

## Results

### Regulation of FUS RNA with retained introns 6 and 7 in WT and ALS-FUS cells

We first sought to establish the molecular basis for preferential retention of FUS introns 6 and 7. Maximum Entropy Modelling (MaxEntScan) analysis (Yeo and Burge, 2004) revealed that the sites flanking the retained region (exon 6 donor/intron 6 donor - E6D/I6D and exon 8 acceptor/intron 7 acceptor - E8A/I7A) are weaker than those of most exon-intron junctions in the gene: E6D/I6D had the lowest donor score, and E8A/I7A had the joint lowest acceptor score (Fig. 1a; Supplementary Table S1), providing a possible mechanism for the less efficient splicing of this region.

**Figure 1.**
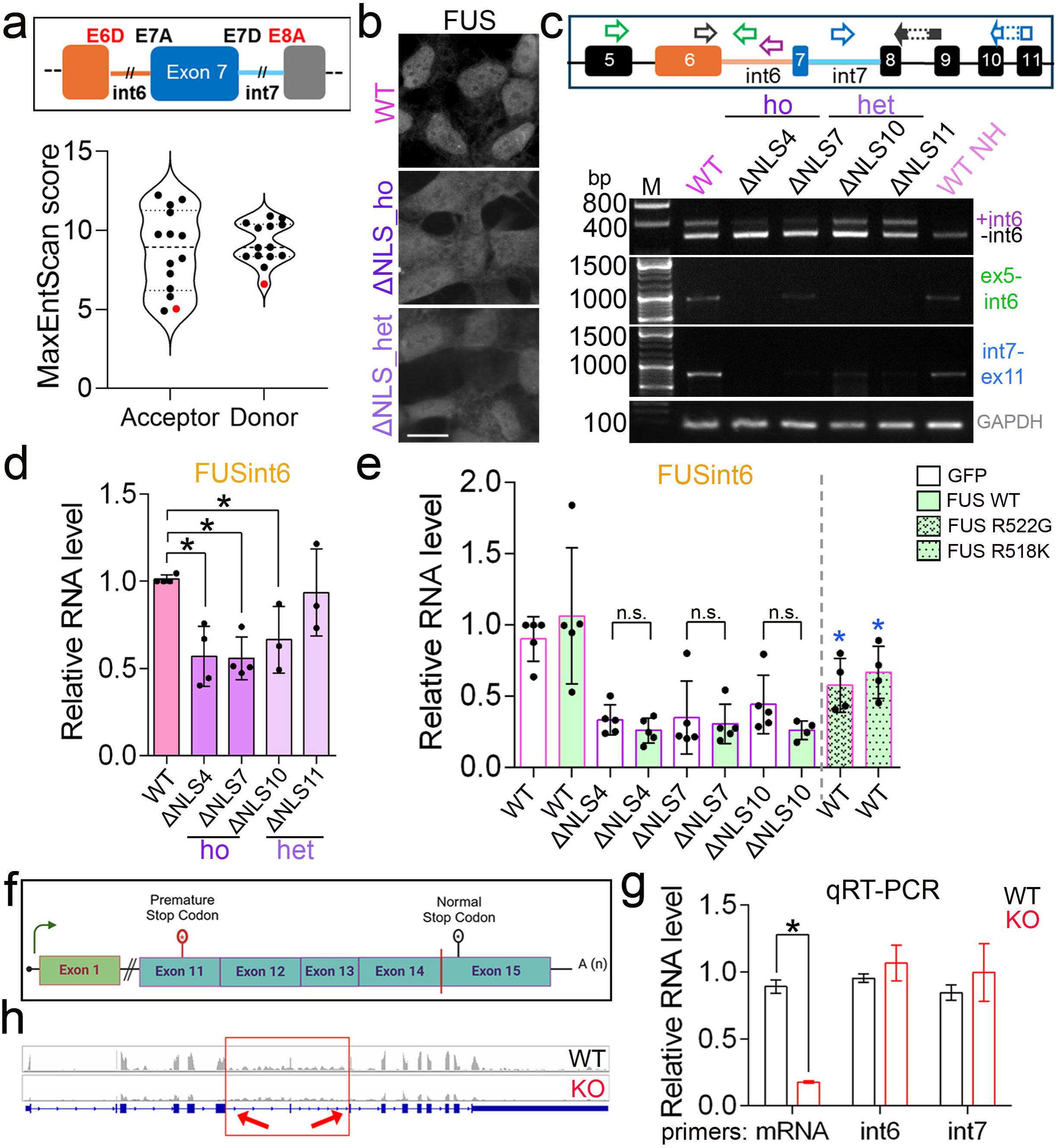
Regulation of FUS RNA with retained introns 6 and 7. **a** Splice sites flanking the retained region in FUSint6&7-RNA are weaker compared to other splice sites in the *FUS* gene. MaxEntScore was used to determine the splice site strength, and the flanking sites (E6D/I6D and E8A/I7) are given in red in the violin plot. **b** FUS mislocalisation in FUSΔNLS cell lines used in the study. **c,d** Reduced retention of introns 6 and 7 in FUS RNA demonstrated by PCR (c) and qRT-PCR (d). WT NH, WT cell samples not heated during RNA extraction. A combination of three primers, FUS_ex6_for, FUS_int6_rev, and FUS_ex8/9_rev, was used to detect intron 6 inclusion. *p<0.05, N=3-4, Kruskal-Wallis with Dunn’s post-hoc test. **e** FUS overexpression does not restore FUS intron 6 retention in FUSΔNLS lines, whereas expression of mutant FUS downregulates FUSint6&7-RNA in WT cells. GFP-tagged FUS WT and ALS-linked variants R522G (mainly cytoplasmic) and R518K (mainly nuclear) were used. N=3-5. *p<0.05, Mann-Whitney *U* test, as compared to WT cells expressing WT FUS-GFP. **f-h** FUSint6&7-RNA level remains unchanged in FUS knockout cells with intact FUS transcription but undetectable FUS protein. *FUS* locus CRISPR-Cas9 editing schematic (f), FUSint6&7-RNA analysis by qRT-PCR (g) and RNAseq (h) are shown. *p<0.05, N=3, Mann-Whitney *U* test.

Consistent with the findings in murine CNS tissue (Humphrey et al., 2020), a reduction in FUSint6&7-RNA was observed in human ALS-FUS cell models – FUSΔNLS SH-SY5Y lines generated by CRISPR-Cas9 editing to endogenously express FUS lacking NLS (An et al., 2019) (Fig. 1b-d). Indeed, ectopic expression of two ALS-FUS mutants, R522G and R518K, significantly downregulated FUSint6&7-RNA (Fig. 1e). However, this reduction could not be rescued by overexpression of GFP-tagged WT FUS (Fig. 1e). This suggested that mutant FUS gain-of-function, rather than a loss-of-function mechanism is responsible for reduced retention of FUS introns 6 and 7 in ALS-FUS models. To address this directly, we used a previously generated FUS knockout (KO) cell line (An et al., 2019) that has a premature stop-codon in the *FUS* gene introduced by CRISPR-Cas9 editing, leading to FUS mRNA degradation by NMD (Fig. 1f). This cell line expresses normal levels of FUS pre-mRNA but has severely reduced FUS mRNA (∼15% of normal) and undetectable protein levels (An et al., 2019) (Fig. 1g). We found that FUSint6&7-RNA levels were not affected in this cell line compared to WT SH-SY5Y cells, as confirmed by qRT-PCR (Fig. 1g) and RNAseq (dataset from An et al., 2019) (Fig. 1h).

These results reveal the importance of a gain-of-function mechanism in affecting the abundance of FUS transcripts with retained introns 6 and 7 in mutant FUS expressing cells.

### FUSint6&7-RNAs form multimolecular foci in the nucleus

Given that the FUSint6&7-RNA species accumulate in the nucleus (Humphrey et al., 2020), we aimed to characterise their fate and regulation in this compartment. We previously showed that a nuclear-localised transcript with retained introns produced from the mouse *Srsf7* gene can form phase-separated granules (Königs et al., 2020). Using an RNA-FISH probe pool covering FUS intron 6 (37 oligonucleotides), bright nuclear foci were detected in all widely used cell lines (HeLa, U2OS, SH-SY5Y, human fibroblasts) (Fig. 2a; Supplementary Figure 1a). In addition to these larger granules, multiple smaller foci were also detectable in the nucleoplasm (Fig. 2a, inset). The smallest/dimmest foci (“1” in Fig. 2a inset) presumably corresponded to single RNA molecules, whereas the intermediate-size foci (“2”) represented their clusters, and the large structures (“3”) likely corresponded to a build-up of FUSint6&7-RNA at the sites of its transcription. Indeed, the number of large foci per cell matched the cell line ploidy (e.g. 3-4 in hypertriploid (3n+) HeLa cells and 2 foci in diploid U2OS and SH-SY5Y cells) (Fig. 2b). Importantly, the large foci were not composed of nascent or uniformly unspliced FUS pre-mRNAs since they were not recognised by a FUS intron 1 specific probe pool (Supplementary Figure 1a). FUSint6&7-RNA were also readily detectable using RNAscope-ISH with chromogenic detection (Fig. 2c). Notably, this detection approach revealed significantly larger granule size in HeLa cells as compared to SH-SY5Y cells, pointing to variable numbers of molecules per focus in different cell lines. Reanalysis of publicly available RNAseq datasets confirmed that a considerable proportion of reads in this region of the *FUS* gene spanned intron 6, exon 7 and intron 7, confirming retention of both introns in the same transcript (data not shown). Consistent with this observation, a FUS intron 7-specific probe pool (33 oligonucleotides) also detected characteristic foci in HeLa cells (Supplementary Figure 1b). In subsequent experiments, either FUSint6 or -int7 specific probes were used to detect endogenous FUSint6&7-RNA foci.

**Figure 2.**
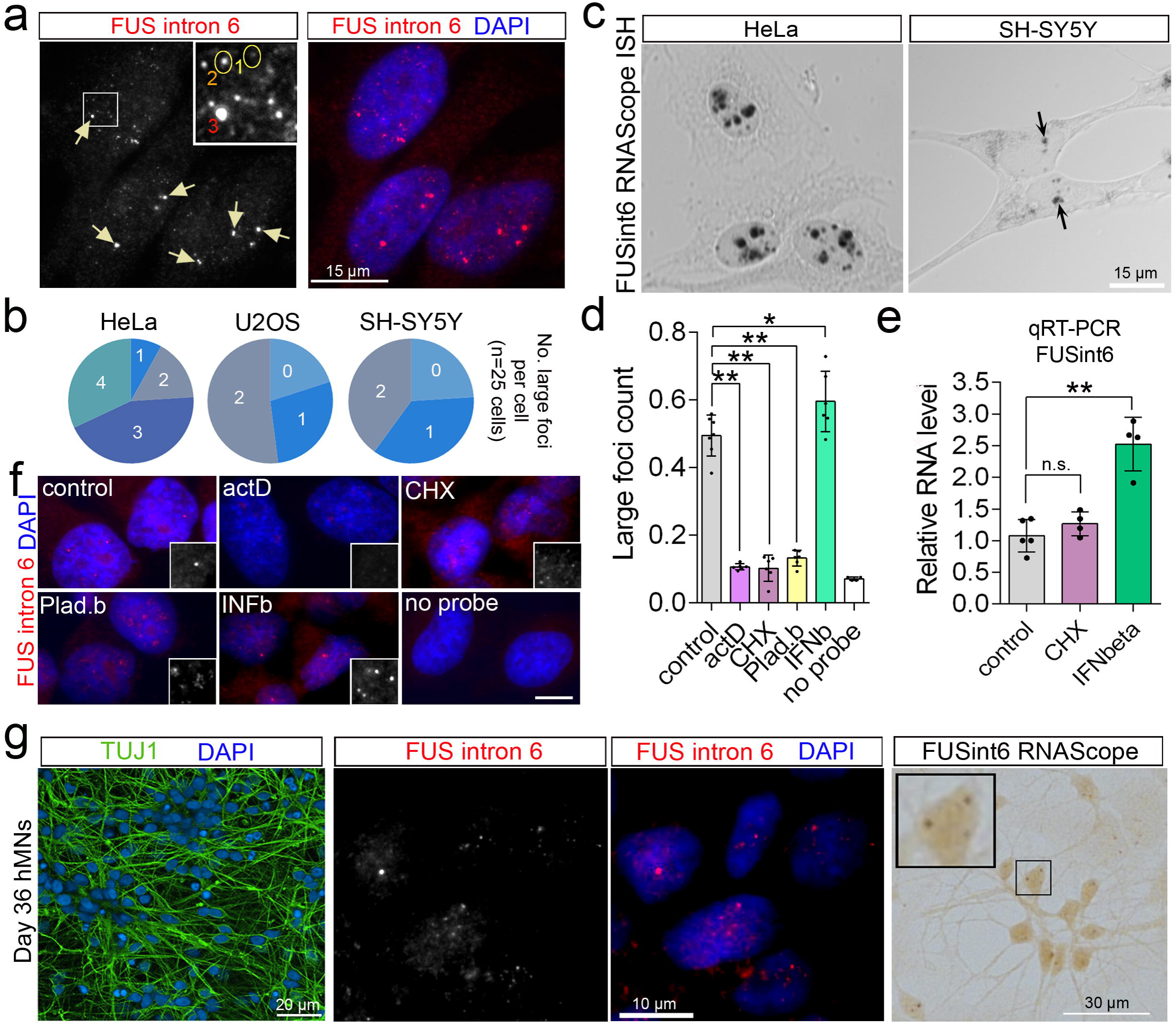
FUS RNA with retained introns forms dynamic nuclear foci. **a** FUSint6&7-RNA forms nuclear foci. Representative images for RNA-FISH with a FUS intron 6 probe in HeLa cells are shown. Three types of foci, based on their size, are labelled. Arrows indicate the large (type “3”) foci, which likely assemble near the sites of transcription. **b** Large foci (type “3”) frequency per cell in different cell lines corresponds to the cell line ploidy. **c** FUSint6&7-RNA foci visualised using FUS intron 6 RNAscope®-ISH probe with chromogenic detection. Representative images for HeLa and SH-SY5Y cells are shown. **d-f** Formation of FUSint6&7-RNA foci relies on ongoing transcription and is sensitive to changes in RNA metabolism. HeLa cells were treated with actinomycin D, CHX or pladienolide B for 4 h, or with IFNbeta for 24 h. FUSint6-positive foci were quantified by an automated high-content imaging assay (d), analysed by qRT-PCR (e) and by high-resolution imaging (f). In d, N=6 (individual wells), *p<0.05, **p<0.01, Kruskal-Wallis with Dunn’s test. In e, N=4-5, **p<0.01, Kruskal-Wallis with Dunn’s test. **g** FUSint6&7-RNA foci form in cultured human motor neurons. Day 36 neurons were used for RNA-FISH and RNAscope®-ISH (FUS intron 6 probes).

Using an automated foci imaging and quantification assay on Opera Phenix high-content imaging system (Supplementary Figure 1c), large FUSint6&7-RNA foci were found to be dispersed by actinomycin D, cychoheximide (CHX) and a splicing inhibitor pladienolide B (Fig. 2d). In contrast, IFNbeta – previously shown to upregulate FUS at the RNA level (Shelkovnikova et al., 2019) – increased the number of foci (Fig. 2d). In line with the previous report (Humphrey et al., 2020), CHX, which is known to inhibit NMD, had no effect on FUSint6&7-RNA abundance, whereas IFNbeta increased it (Fig. 2e). Intriguingly, CHX and pladienolide B induced fragmentation of the large foci (type “3”) into smaller dots without changes to the total signal intensity (Fig. 2f). This suggested that factors other than FUSint6&7-RNA itself are required for maintaining the foci integrity.

Finally, we confirmed that nuclear FUSint6&7-RNA granules form in an ALS-relevant cell type, human motor neurons (Fig. 2g). Similar to non-neuronal cells, FUSint6&7-RNA expression remained unchanged in response to CHX and was upregulated by IFNbeta (Supplementary Figure 1d).

Thus, partially processed FUS transcripts can assemble dynamic multimolecular structures in the nuclei of diverse cell types.

### FUSint6&7-RNA foci are speckle-interacting condensates scaffolded by intron 7

We next focused on detailed characterisation of the FUSint6&7-RNA foci. Both large and small foci did not overlap with known nuclear bodies but were typically (>80% of population) present on the border of, or “docked” to, nuclear speckles (Fig. 3a and not shown). This pattern resembled that of paraspeckles, which are nuclear bodies assembled by lncRNA NEAT1_2, that both cluster at the transcription site and distribute throughout the nucleus, becoming associated with speckles (Takakuwa et al., 2023). Notably, RNase A treatment in semi-permeabilised cells eliminated the large FUSint6&7-RNA foci, whereas DNase I treatment had no effect (Fig. 3b).

**Figure 3.**
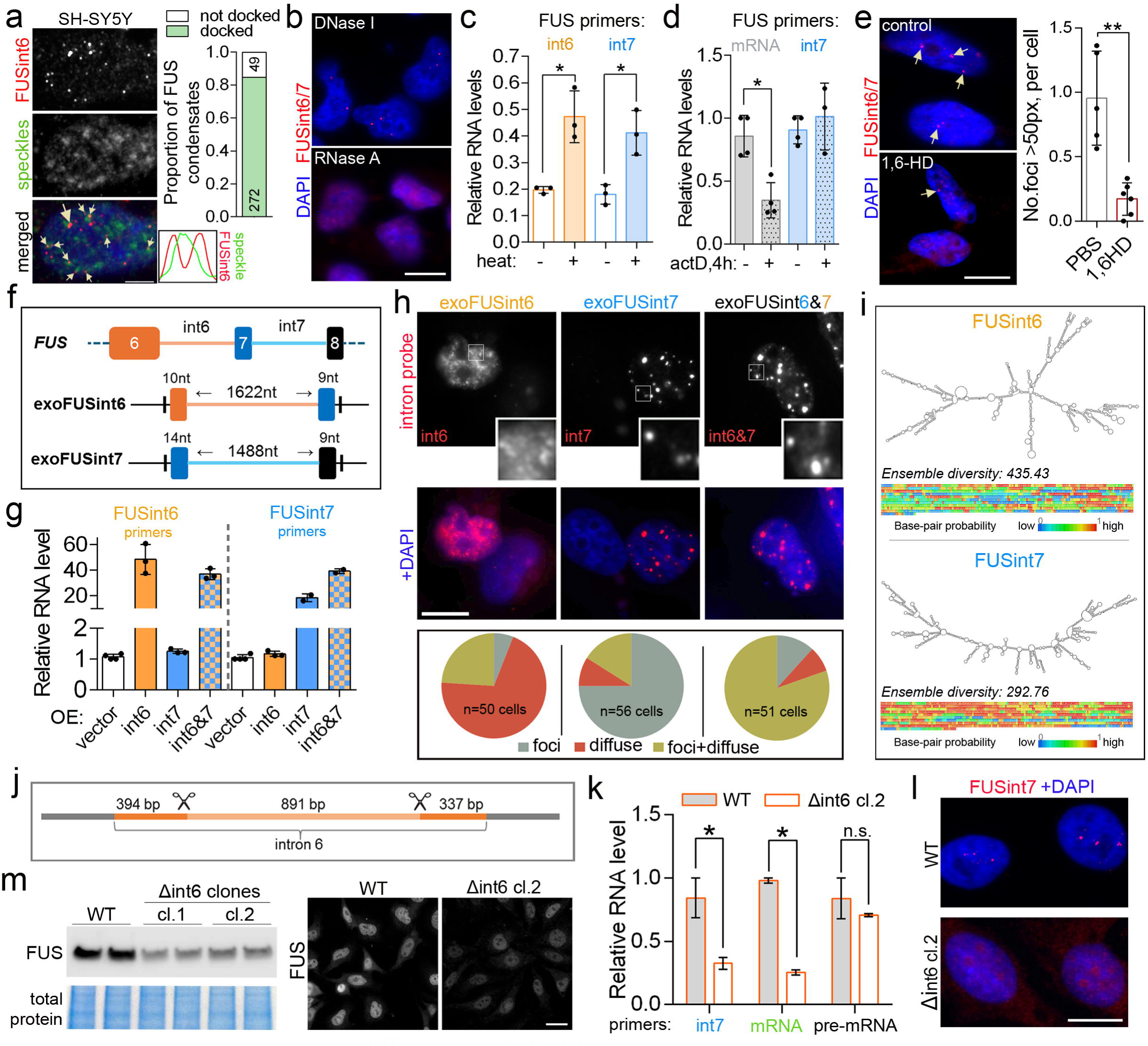
FUSint6&7-RNA foci are phase-separated condensates nucleated by intron 7 and associated with splicing speckles. **a** FUSint6&7-RNA foci are associated with splicing speckles. Proportion of foci physically interacting with speckles (“docked”) was quantified for each cell. Numbers of individual condensates of each type are indicated within bars (N=20 cells). SH-SY5Y cells that possess prominent speckles were used, and a representative image is shown. PNN was used to visualise speckles. Scale bar, 2 μm. **b** RNase A treatment eliminates FUSint6&7-RNA foci. Semi-permeabilised HeLa cells were treated with RNase A or DNase I. Representative images are shown. Scale bar, 10 μm. **c** FUSint6&7-RNA is semi-extractable. qRT-PCR analysis (FUS intron 6- or 7-specific primers) in samples with or without heating (55⁰C for 10 min) prior to RNA isolation. HeLa cells were used. N=3, *p<0.05, Mann-Whitney *U* test. **d** FUSint6&7-RNA has a high stability. FUSint6&7-RNA (intron 6 primers) and FUS mRNA levels were analysed by qRT-PCR in HeLa cells after a 4-h actinomycin D treatment. GAPDH was used for normalisation. N=3-4, *p<0.05, Mann-Whitney *U* test. **e** FUSint6&7-RNA foci are sensitive to a LLPS-disrupting agent. 1,6-hexanediol (1,6-HD) treatment was performed in semi-permeabilised HeLa cells. Representative images and quantification are shown. N=5-6 (fields of view, FoV), analysed in a representative experiment; **p<0.01, Mann-Whitney *U* test. Scale bar, 10 μm. **f,g** Efficient and comparable expression of exogenous FUS introns 6 and 7 upon transient transfection of the intronic sequences. Schematic of the constructs for ectopic expression of FUS introns (f) and qRT-PCR analysis using intron 6- or 7-specific primers (g) are shown. N=3-4. **h** FUS intron 7, but not intron 6, forms compact, dense nuclear condensates upon ectopic expression. Representative images and quantification of phenotypes are shown. Scale bar, 5 μm. **i** FUS intron 7 is predicted to be more structured than intron 6, with lower ensemble diversity. Minimum free energy (mfe) structure, alongside base-pairing probabilities heatmaps, for FUS introns 6 and 7, was generated by RNAfold. **j-m** CRISPR-mediated deletion of a middle portion of FUS intron 6 destabilises FUSint6&7-RNA and leads to FUS mRNA and protein depletion. Positions of sgRNAs used (j), levels of FUS RNA species (k), FUSint6&7-RNA condensate analysis (l) and FUS protein levels (m) in homozygous Δint6 HeLa clones are shown. In *l* and *m*, representative images are shown. N=3-4, *p<0.05, Mann-Whitney *U* test, n.s., non-significant. Scale bars, 10 μm in *l* and 20 μm in *m*.

Architectural RNAs (arcRNAs) that assemble phase-separated nuclear bodies are tightly packed in RNPs, such that heating or mechanical shearing is required for their efficient isolation during RNA purification with a Trizol-type reagent (Chujo et al., 2016; Chujo et al., 2017). Indeed, FUSint6&7-RNA had semi-extractable properties; when the heating step was omitted (included as a standard into our RNA isolation protocol, see Materials and Methods), the yield of this RNA was significantly decreased (Fig. 3c). Although transcription block prevents granule formation by arcRNAs (Chujo et al., 2016), the RNA itself can remain stable for hours, for example, as in the case of NEAT1_2 (Clark et al., 2012). We found that FUSint6&7-RNA level remains unchanged after a 4-h actinomycin D treatment (when normalised to GAPDH that has a half-life of ∼8 h, Dani et al., 1984), whereas FUS mRNA level was decreased to ∼25% at this time-point (Fig. 3d). Condensates formed by liquid-liquid phase separation (LLPS) are sensitive to 1,6-hexanediol (1,6-HD) – an aliphatic alcohol that disrupts weak electrostatic interactions (Jain et al., 2016). 1,6-HD treatment in semi-permeabilised cells indeed decreased the number of FUSint6&7-RNA foci (Fig. 3e).

We next examined the contribution of introns 6 and 7 to FUSint6&7-RNA condensate assembly. Constructs for ectopic expression of the two introns (exoFUSint6 and -7 thereafter) were generated, by cloning the full intronic sequence with the splice sites into an expression vector (Fig. 3f). qRT-PCR confirmed accumulation of both introns when expressed separately or co-expressed (Fig. 3g). RNA-FISH and nuclear-cytoplasmic fractionation demonstrated that although both overexpressed introns were largely retained in the nucleus, exoFUSint7 readily formed dense foci, whereas exoFUSint6 remained largely diffuse (albeit we could not exclude the formation of small condensates masked by the diffuse signal) (Fig. 3h; Supplementary Figure 2a). FUS intron 1, which has a similar length (2170 nt) but is not retained, also remained diffuse when overexpressed (Supplementary Figure 2b). Nuclear foci formed by exoFUSint7 did not overlap with known nuclear bodies – paraspeckles (NEAT1_2), Cajal bodies (coilin p80) or Gems (SMN) (Supplementary Figure 2c). Furthermore, exoFUSint7 condensates responded to CHX and pladienolide B treatments in a way similar to their endogenous counterparts (Supplementary Figure 2d). RNAfold predictions (Gruber et al., 2008) indicated that intron 7 is more structured as compared to intron 6, with lower positional entropy, higher probability of base pairing and lower ensemble diversity (Fig. 3i).

To investigate the sequence requirements for FUSint6&7-RNA condensate assembly, we targeted FUS introns 6 and 7 using specific CRISPR-Cas9 sgRNAs. While we were unable to obtain viable clones lacking intron 7 sequences, we successfully generated HeLa cells with a large portion of intron 6 deleted – Δint6 clones (Fig. 3j, Supplementary Figure 3). FUSint6&7-RNA (measured with an intron 7-specific primer pair) was significantly downregulated in these cell clones (Fig. 3k). This was accompanied by the loss of FUSint6&7-RNA condensates (Fig. 3l), reduced expression of the FUS mRNA (Fig. 3k) and FUS protein (Fig. 3m). Notably, the mutation had no detectable effect on *FUS* transcriptional activity as measured by qRT-PCR with intron 1-specific primers (Fig. 3k). We also confirmed that the *FUS* copy number was not affected by CRISPR-Cas9 editing (data not shown).

These data indicate that the foci containing multiple copies of FUSint6&7-RNA exhibit properties of biomolecular condensates. Their assembly in the vicinity of nuclear speckles appears to depend on the well-structured intron 7 and LLPS. Intron 6, on the other hand, may be required for proper processing or/and stability of FUS transcripts.

### FUSint6&7-RNA condensates are positive regulators of FUS expression

Since FUS intron 7 in isolation can assemble condensates, we hypothesised that over-expressing intron 7 sequences may disrupt endogenous FUSint6&7-RNA condensates due to sequestration of their essential components. To test this prediction, we used a combination of a FUSint6-specific RNA probe pool (Stellaris) – for detection the endogenous FUSint6&7-RNA, and a single FUSint7-specific oligonucleotide probe – for detection of exoFUSint7 condensates. This FUSint7 probe had low labelling efficiency and detected exoFUSint7 condensates but not endogenous FUSint6&7-RNA condensates (Fig. 4a). FUSint6&7-RNA condensates were found dissolved in the majority of cells that developed exoFUSint7 *de novo* condensates (Fig. 4a). Interestingly, we also observed fusion events between the endogenous and exogenous condensates (Fig. 4a, inset).

**Figure 4.**
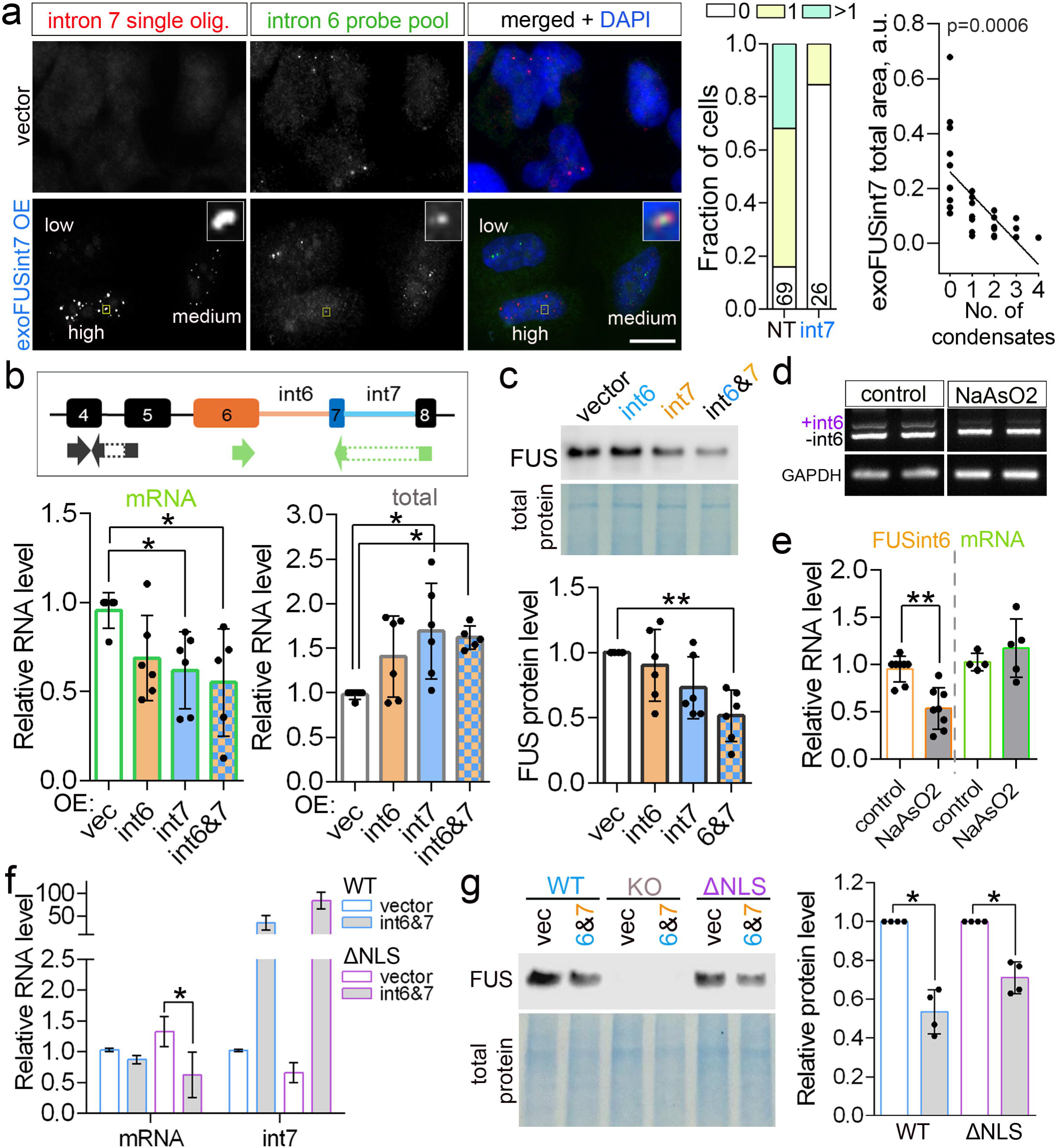
FUSint6&7-RNA condensates regulate FUS mRNA levels. **a** Ectopic expression of FUS intron 7 dissolves endogenous FUSint6&7-RNA condensates. Single FUSint7 oligonucleotide probe recognising exoFUSint7 condensates but not endogenous FUSint6&7-RNA condensates was used in combination with the Stellaris FUSint6 probe pool. Representative images and quantification of endogenous FUSint6&7-RNA condensates are shown. Cells with high, medium and low exoFUSint7 expression are indicated. Inset shows fusion of a FUSint6&7-RNA condensate with an exoFUSint7 condensate. Number of endogenous condensates quantified in exoFUSint7 condensate-containing cells (int7) *vs.* non-transfected cells (NT) in the same FoV is indicated inside the bars. Correlation between the area of exoFUSint7 signal in individual nuclei and the number of endogenous FUSint6&7-RNA condensates are also shown (N=25 cells). **b** Ectopic expression of exoFUSint7 or exoFUSint6+7 leads to FUS mRNA downregulation. qRT-PCR analysis of FUS total (FUS mRNA + FUSint6&7-RNA) and mRNA levels is shown. N=3-5, *p<0.05, Kruskal-Wallis with Dunn’s test. **c** Ectopic expression of exoFUSint6+7 leads to FUS protein downregulation. Representative western blot and quantification for HeLa cells are shown. N=6, **p<0.01, Kruskal-Wallis with Dunn’s test. **d,e** FUS intron retention is responsive to cellular stress. FUS RNA levels were analysed in cells recovering from NaAsO2 stress (1 h stress+3 h recovery) by PCR with a triple primer combination (d) and qRT-PCR (e). N=5-7, **p<0.01, Mann-Whitney *U* test. **f,g** exoFUSint7 expression downregulates FUS mRNA and protein in an ALS-FUS cell model. FUSΔNLS lines were analysed by qRT-PCR (f) and western blot (g). N=3-4, *p<0.05, Mann-Whitney *U* test. In *f*, ΔNLS10 was used and in *g*, ΔNLS4 was used.

Analysis of FUS mRNA levels in cells expressing exoFUSint7, separately or in combination with exoFUSint6, demonstrated that these treatments triggered a significant downregulation of FUS mRNA (by ∼30%; Fig. 4b). This was accompanied by an increase in the total FUS transcript levels (FUS mRNA+FUSint6&7-RNA) (Fig. 4b), due to upregulation of endogenous FUSint6&7-RNA (detected using intron 6-specific primers) (Supplementary Figure 4a,b). Interestingly, exoFUSint6 alone also upregulated FUSint6&7-RNA (Supplementary Figure 4a), however it did not significantly change the abundance of FUS mRNA (Fig. 4b). This milder effect was consistent with the limited condensate-forming ability of intron 6. Importantly, FUS protein level was decreased in exoFUSint6+7 expressing cells, mirroring mRNA downregulation (Fig. 4c). FUS intron 1 overexpression did not affect FUS mRNA levels (Supplementary Figure 4c). This data suggested that FUSint6&7-RNA condensates may represent reservoirs of FUS transcripts poised for the production of FUS mRNA, whose disruption has a suppressive effect on the expression of this gene.

To further test the reservoir model, we analysed FUS expression under cellular stress conditions. Stress treatments have been reported to cause global mRNA decay (Bresson et al., 2020; Dar et al., 2024). We found that FUSint6&7-RNA and its condensates become depleted during the recovery from sodium arsenite stress, as well as in response to other chemical stresses (Fig. 4d,e; Supplementary Figure 4d,e). This was associated with maintained FUS mRNA level (Fig. 4d,e), without significant changes in *FUS* transcription (pre-mRNA level, Supplementary Figure 4f).

Finally, we showed that FUS mRNA and protein were downregulated following exoFUSint7 overexpression in FUSΔNLS neuroblastoma cells (Fig. 4f,g). Of note, exoFUSint6/7 expression did not cause significant cellular toxicity (Supplementary Figure 4g).

Thus, the higher-order assemblies of FUSint6&7-RNA appear to be an integral part of normal FUS mRNA production, contributing to the regulation of this process in response to external cues.

### FUSint6&7-RNA condensates accumulate splicing factors and the m6A reader YTHDC1

To gain insights into FUSint6&7-RNA condensate composition and hence regulation, we performed hybridization-proximity labelling coupled with mass-spectrometry (HyPro-MS) (Yap et al., 2022a, b) (Fig. 5a). We previously used this approach to analyse protein and RNA interactomes of RNA-seeded compartments in genetically unperturbed cells. Here, we leveraged an enhanced version of the HyPro procedure, which allows efficient labelling of small RNP structures while reducing the unspecific diffusion of activated biotin (*Yap et al., submitted*). A probe set covering FUS introns 6 and 7 (68 oligonucleotides in total) was used, alongside two controls: a no-probe control (Ctrl) and beta-actin (ACTB) intronic probe set (34 oligonucleotides). The latter probe detected small foci in HeLa cells corresponding to newly produced pre-mRNAs (Supplementary Figure 5a) – and therefore was ideally suited as a control for the analysis of FUSint6&7-RNA specific interactors. Efficient labelling of FUSint6&7-RNA condensates in a HyPro experiment was confirmed by combining HyPro-FISH (Yap et al., 2022b) with FUSint6&7-RNA specific probes and RNA-FISH with the FUS intron 6 Stellaris probe set (Fig. 5b).

**Figure 5.**
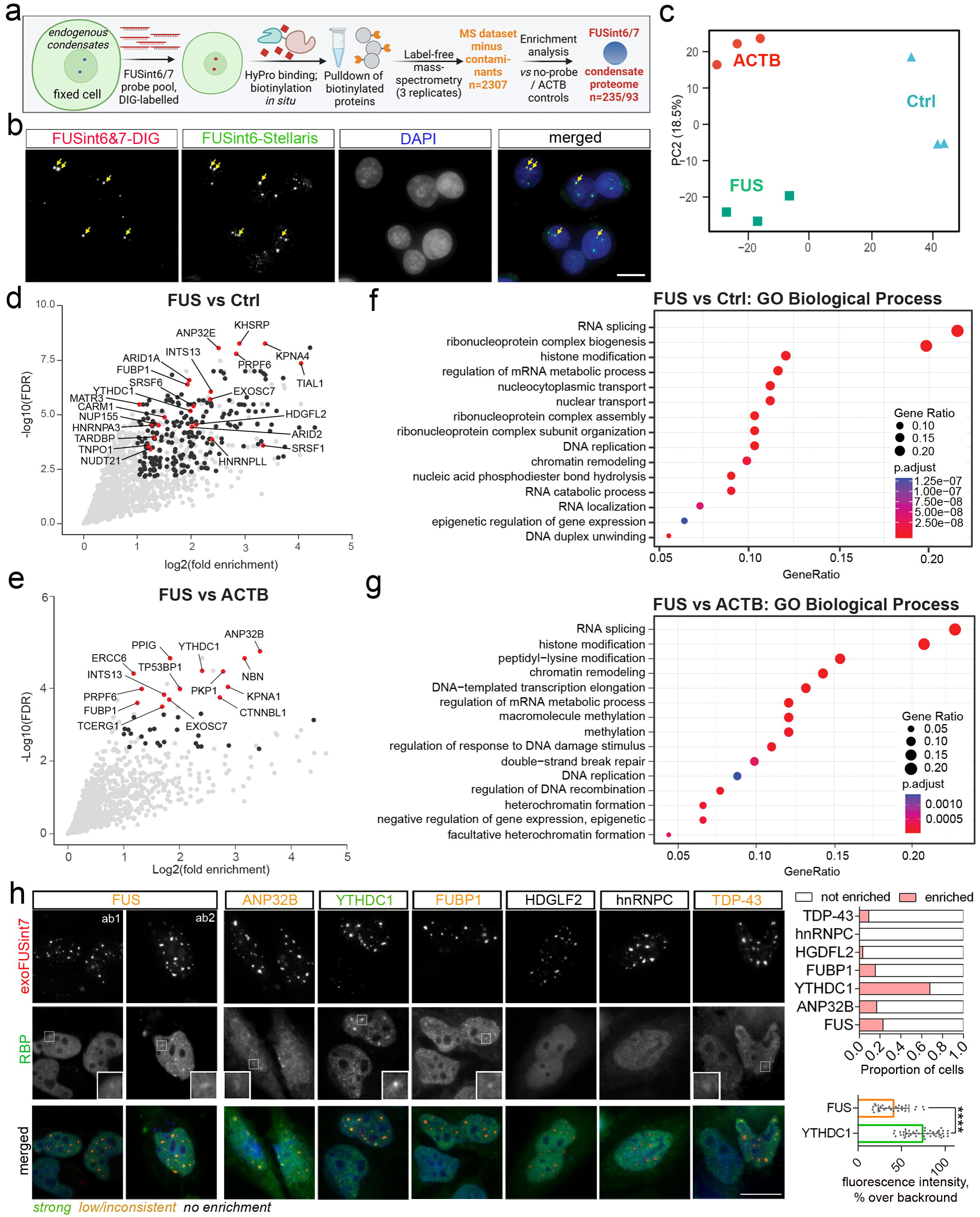
Proteomic analysis of FUSint6&7-RNA condensates by enhanced HyPro-MS. **a** Experimental pipeline for HyPro-MS analysis of FUSint6&7-RNA condensates. **b** Efficient labelling of the endogenous FUSint6&7-RNA condensates using HyPro probes. Stellaris FUSint6-specific probe was used for co-staining. Arrows indicate the foci labelled by both Stellaris and HyPro probes. Scale bar, 10 µm. **c** Principal component analysis (PCA) demonstrating clustering of triplicated HyPro-MS samples for both probes and the no-probe control. **d** Volcano plot for FUSint6&7-RNA condensates versus no-probe control (Ctrl). Proteins with padj<0.05 are labelled in black. Proteins with padj<0.05 and involved in RNA metabolism-related pathways and/or implicated in neurodegeneration, are labelled in red. **e** Volcano plot for FUSint6&7-RNA condensates versus ACTB probe. Proteins with padj<0.1 are labelled in black and the top 15 hits are labelled in red. **f** Dot plot of GO Biological Process term enrichment analysis for nuclear proteins identified in FUSint6&7-RNA condensates and significantly enriched as compared to no-probe control. **g** Dot plot of GO Biological Process term enrichment analysis for nuclear proteins identified in FUSint6&7-RNA condensates and significantly enriched as compared to ACTB probe control. **h** Validation of HyPro-MS proteins enriched in FUSint6&7-RNA condensates, as well as FUS and other proteins previously shown to bind FUSint6&7-RNA. Cells expressing exoFUSint7 were analysed by RNA-FISH and immunofluorescence with appropriate antibodies. Representative images are shown. Top graph, 28-35 transfected cells with condensates were analysed per protein; bottom graph, 41 and 27 individual condensates were analysed for YTHDC1 and FUS enrichment, respectively, ****p<0.0001, two-tailed unpaired *t* test.

Following the hybridization with DIG-labelled probes and incubation with the HyPro enzyme (see Materials and Methods for details), the molecules physically proximal to the FUSint6&7-RNA transcripts (bait) were biotinylated *in situ* and captured on streptavidin beads under denaturing conditions. The expected enrichment of biotinylated RNA “baits” on the beads was confirmed by qRT-PCR (Supplementary Figure 5b). Efficient protein biotinylation in all samples was confirmed by western blotting (Supplementary Figure 5c). Label-free mass-spectrometry of the captured proteins identified a total of 2869 proteins – reduced to 2307 after additional filtering steps (removal of common contaminants, including those from the CRAPome database; https://reprint-apms.org/). Principal component analysis (PCA) of the filtered proteins demonstrated good clustering of the triplicates for each condition (Fig. 5c).

Enrichment analysis for FUSint6&7-RNA condensates as compared to the no-probe control (Ctrl) identified 235 proteins (p<0.05), of which 184 were significant after correction for multiple comparisons (padj<0.05) (Supplementary Table S2; Fig. 5d). Only 93 proteins were significantly enriched when compared to ACTB probe (p<0.05), of which 19 remained significant after correction for multiple comparisons (padj<0.05) (Supplementary Table S2; Fig. 5e). FUSint6&7-RNA condensates were enriched in factors involved in RNA splicing, histone modification and RNP complex assembly (Fig. 5f). When compared to ACTB intronic foci, they also showed enrichment for the factors related to macromolecule methylation and DNA damage repair (Fig. 5g). The top ten hits (vs. ACTB probe) included nuclear speckle components such as SR proteins, PPIG (Lin et al., 2004), and the nuclear N6-methyladenosine (m6A) reader YTHDC1 known to be enriched in speckles (Wang et al., 2021). These condensates were also found to recruit ALS-linked proteins TDP-43, MATR3, EWS and hnRNPA3 (Fig. 5d,e; Supplementary Table S2).

Given the importance of FUS intron 7 for condensate assembly established in the above experiments (Fig. 3), we used exoFUSint7 for immunofluorescence validation of the HyPro-MS hits. In addition to the HyPro-MS top hits ANP32B, YTHDC1, FUBP1, PPIG, TCERG1, we included proteins with known binding motifs in FUS introns 6/7 (RBPDB database, http://rbpdb.ccbr.utoronto.ca/) such as HDGFL2 and hnRNPC, as well as TDP-43 and FUS itself. YTHDC1 was found to be highly enriched in >60% of all exoFUSint7 condensates, and TDP-43, ANP32B and FUBP1 also showed some enrichment in a fraction (<20%) of condensates (Fig. 5h). Other protein candidates showed no detectable enrichment (Fig. 5h and data not shown), indicating that they may interact with the sequences outside of intron 7 or associate with the condensates only transiently. FUS protein, which showed only minimal enrichment in the HyPro-MS analysis using ACTB as a control (1.05-fold logFC vs. ACTB, p=0.044, padj=0.23), displayed some recruitment into ∼20% of exoFUSint7 condensates; however, this recruitment was significantly weaker compared to YTHDC1 (Fig. 5h).

Thus, FUSint6&7-RNA condensates interact with a distinct set of proteins involved in post-transcriptional regulation of gene expression.

### m6A and YTHDC1 promote FUSint6&7-RNA condensation

The m6A reader YTHDC1 was strongly enriched in FUS RNA condensates according to both HyPro-MS (top 5 enriched proteins vs. ACTB probe; FC=2.41, padj=0.011) and immunofluorescence validation (Fig. 5h). Remarkably, YTHDC1 knockdown dissolved both large and small condensates (Fig. 6a,b; Supplementary Figure 6a), without altering total FUSint6&7-RNA level (Fig. 6c). However, FUS mRNA was downregulated following YTHDC1 knockdown (Fig. 6c), in line with the contribution of the condensates to post-transcriptional splicing of FUSint6&7-RNA.

**Figure 6.**
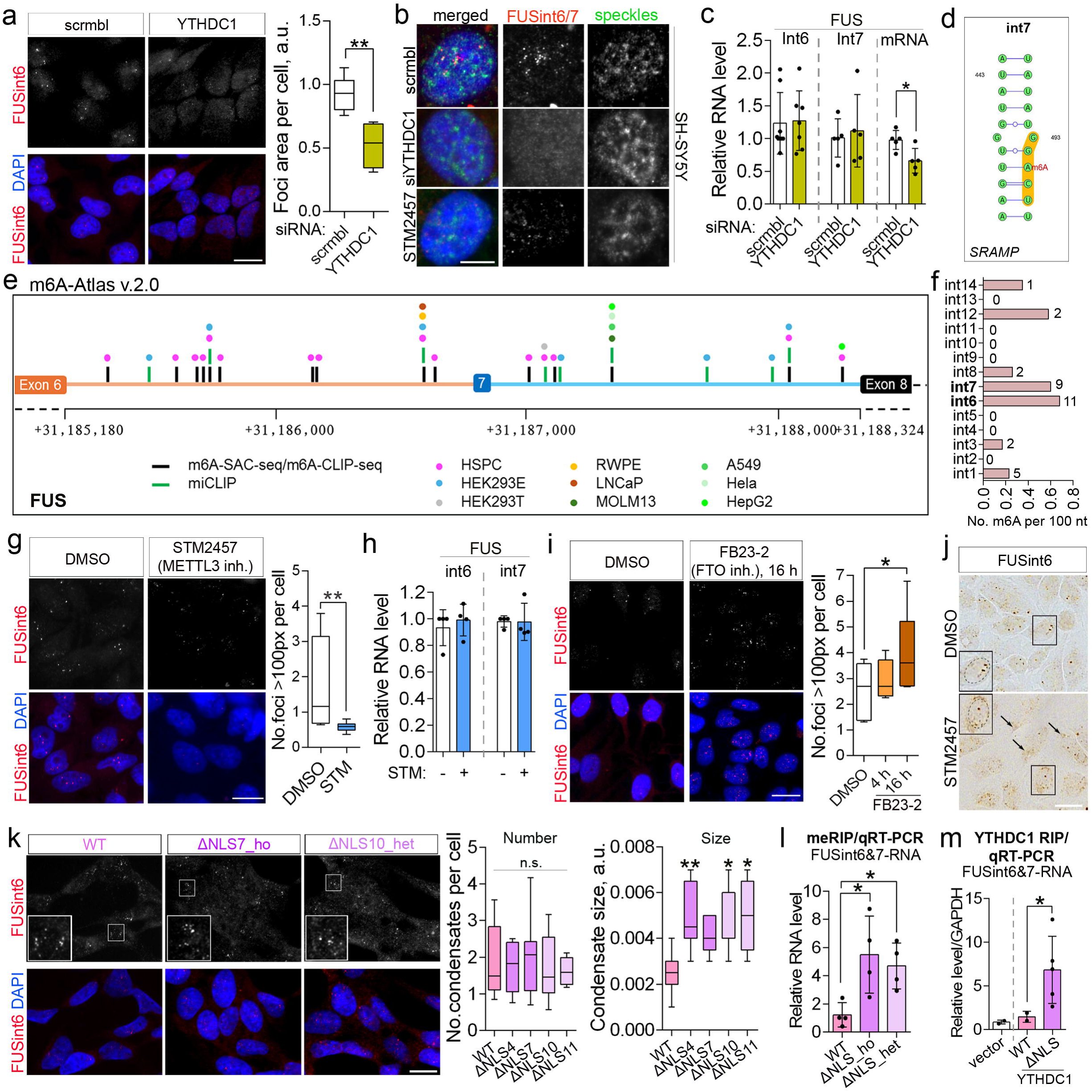
m6A/YTHDC1 maintain FUSint6&7-RNA condensates. **a,b** YTHDC1 depletion or m6A downregulation leads to a loss of FUSint6&7-RNA condensate integrity. Representative images (general plane) and quantification of condensates (a) as well as high-resolution images (b) are shown. 107 and 65 cells (4 or 5 FoV) were analysed for scrambled and YTHDC1 siRNA, respectively, from a representative experiment. **p<0.01, Mann-Whitney *U* test. HeLa cells were used in *a* and SH-SY5Y cells were used in *b*. Representative images are shown. STM2457-treated cells are also shown. In *b*, PNN was used as a speckle marker. Scale bar, 20 μm in *a* and 5 μm in *b*. **c** YTHDC1 depletion does not affect FUSint6&7-RNA level but downregulates FUS mRNA. qRT-PCR analysis with FUS intron 6- and 7-specific primers was performed in HeLa cells. N=5-7, *p<0.05, Mann-Whitney *U* test. **d** High-confidence DRACH motif in FUS intron 7 shown in a structural context, as predicted by SRAMP. **e,f** FUS introns 6 and 7 are extensively methylated. m6A-Atlas 2.0 database was used for mapping methylation sites (e) and calculating the m6A mark density (f). Also see Supplementary Table S3. **g,h** Pharmacological inhibition of a m6A writer METTL3 affects FUSint6&7-RNA condensate integrity without changing levels of this RNA. Representative images and quantification of the condensate number in STM2457-treated cells (g) and qRT-PCR analysis of RNA level (h) are shown. In *g*, cells were treated for 16 h, and >200 cells (5 FoV) were analysed per condition from a representative experiment, **p<0.01, Mann-Whitney *U* test. Scale bar, 20 μm. In *h*, qRT-PCR analysis was done using FUS intron 6- and -7 specific primers. N=4. **i** Pharmacological inhibition of a m6A eraser FTO promotes FUSint6&7-RNA condensate assembly. Representative images and quantification of the condensate number in FB23-2 treated cells are shown. Cells were treated with the compound for either 4 or 16 h. 79, 106 and 117 cells (5 FoV) were analysed for DMSO, 4-h FB23-2 and 16-h FB23-2 treatments, respectively, from a representative experiment. *p<0.05, Kruskal-Wallis with Dunn’s test. Scale bar, 20 μm. **j** FUS intron 6 RNAscope-ISH reveals partial cytoplasmic redistribution of FUSint6&7-RNA in STM2457-treated cells. Arrows indicate cytoplasmic accumulations of this RNA; nucleus is circled in the insets. Scale bar, 20 μm. **k** FUSΔNLS lines have preserved or enhanced ability to form FUSint6&7-RNA condensates. Representative images and quantification are shown. 234, 239, 234, 141 and 62 cells were analysed for WT, ΔNLS4, ΔNLS7, ΔNLS10 and ΔNLS11 lines, respectively (from 4-9 FoV). *p<0.05, **p<0.01, Kruskal-Wallis with Dunn’s test. **l** MeRIP demonstrates increased FUSint6&7-RNA methylation in FUSΔNLS lines. Total RNA purified from ΔNLS7 (homozygous) and ΔNLS10 (heterozygous) lines was subjected to pulldown using m6A antibody-coated beads and used for qRT-PCR analysis with FUSint6-specific primers. Methylated FUSint6&7-RNA level was normalised to GAPDH mRNA (extensively methylated transcript), then to total FUSint6&7-RNA level in the respective cell line, and finally, to no-antibody beads control. N=4, *p<0.05, Kruskal-Wallis with Dunn’s test. **m** Enhanced association of FUSint6&7-RNA with YTHDC1 in FUSΔNLS lines. RIP was performed using Flag-Trap agarose in YTHDC1-Flag expressing WT and FUSΔNLS cells, followed by qRT-PCR analysis with FUSint6-specific primers. Venus-Flag was used as a control (“vector”). FUSint6&7-RNA level was normalised to GAPDH and then to total FUSint6&7-RNA level in the respective cell line. Results for ΔNLS4, ΔNLS7 and ΔNLS10 lines were combined. N=2. *p<0.05, Mann-Whitney *U* test (WT *vs*. ΔNLS).

YTHDC1 enrichment suggested that either FUSint6&7-RNA itself or other RNAs in these condensates were methylated. Both FUS intron 6 and 7 contain several copies of the m6A modification consensus motif DRACH, one of which (GGACU in intron 7) is a predominant m6A-modified motif and a predicted high-confidence m6A site in this RNA by SRAMP (Zhou et al., 2016) (Fig. 6d). Intron 7 also contains a YTHDC1 binding motif GAAUGC (http://rbpdb.ccbr.utoronto.ca/) (Cook et al., 2011). Analysis of public datasets via the m6A-Atlas 2.0 aggregator confirmed multiple m6A marks in both introns in the majority of cell lines (11 and 9 in introns 6 and 7, respectively) (Fig. 6e; Supplementary Table S3). When normalised to intron length, these two introns were found to harbour more m6A marks than other *FUS* introns (Fig. 6f; Supplementary Table S3). To address whether methylation levels modulate FUSint6&7-RNA condensates, we used STM2457, a specific inhibitor of a major m6A writer/methyltransferase, METTL3 (Yankova et al., 2021). Similar to YTHDC1 knockdown, pharmacological depletion of the m6A mark impacted the condensate integrity without affecting FUSint6&7-RNA level (Fig. 6b,g,h). This result was corroborated by siRNA-mediated depletion of another m6A writer, METTL16 (Supplementary Figure 6b). Conversely, m6A upregulation by FB23-2, a small molecule inhibitor of an m6A eraser FTO (Huang et al., 2019), increased the condensate abundance (Fig. 6i). FUSint6 RNAScope-ISH also revealed cytoplasmic redistribution of FUSint6&7-RNA in STM2457-treated cells (Fig. 6j), indicating that loss of condensate integrity is associated with reduced nuclear retention.

Surprisingly, the numbers of FUSint6&7-RNA condensates remained unchanged, and their size increased in the FUSΔNLS neuroblastoma cell lines (Fig. 6k), despite FUSint6&7-RNA downregulation (Fig. 1; heated samples were used for qRT-PCR analysis, ruling out the contribution of semi-extractability). We hypothesised that this effect could be due to increased condensation of FUSint6&7-RNA in response to elevated m6A methylation. Indeed, m6A RNA immunoprecipitation (MeRIP) coupled with qRT-PCR revealed a significant gain in m6A modification on FUSint6&7-RNA in these cell lines (adjusted for the total transcript level; Fig. 6l). To corroborate this result, we performed RNA immunoprecipitation (RIP) in WT and FUSΔNLS cells that transiently expressed YTHDC1-Flag protein, and measured FUSint6&7-RNA levels by qRT-PCR. MALAT1, which is extensively m6A-modified (Wang et al., 2021), demonstrated ∼40-fold enrichment in the YTHDC1-Flag IP samples, as compared to the vector-only control, confirming successful pulldown (Supplementary Figure 6c). FUSint6&7-RNA was dramatically enriched in YTHDC1-Flag IP preparations from FUSΔNLS cells as compared to WT cells (Fig. 6m). Of note, some downregulation of YTHDC1 mRNA was detectable by RNAseq in the heterozygous FUSΔNLS lines (Supplementary Figure 6d); however, the abundance and distribution of YTHDC1 protein remained unchanged (Supplementary Figure 6e). Global m6A levels were also unaltered in these cell lines, as indicated by dot-blot and ELISA analyses (Supplementary Figure 6f,g). We concluded that mutant FUS drives an increase in m6A modification of FUSint6&7-RNA, which in turn promotes condensation of this RNA in the nucleus.

### m6A modification limits FUS protein association with FUSint6&7-RNA

FUS protein has multiple binding sites in FUS introns 6 and 7 according to iCLIP (Humphrey et al., 2020; Rogelj et al., 2012), yet we observed only modest FUS protein recruitment into FUSint6&7-RNA condensates in HyPro-MS and immunostaining validation studies (Fig. 5). Ectopically expressed FUS-GFP also did not display detectable enrichment in endogenous FUSint6&7-RNA condensates and had no effect on the condensate size (Fig. 7a,b). Since m6A can both attract and repel RBPs (Edupuganti et al., 2017), we hypothesised that this modification on FUSint6&7-RNA may regulate FUS protein association with these condensates.

**Figure 7.**
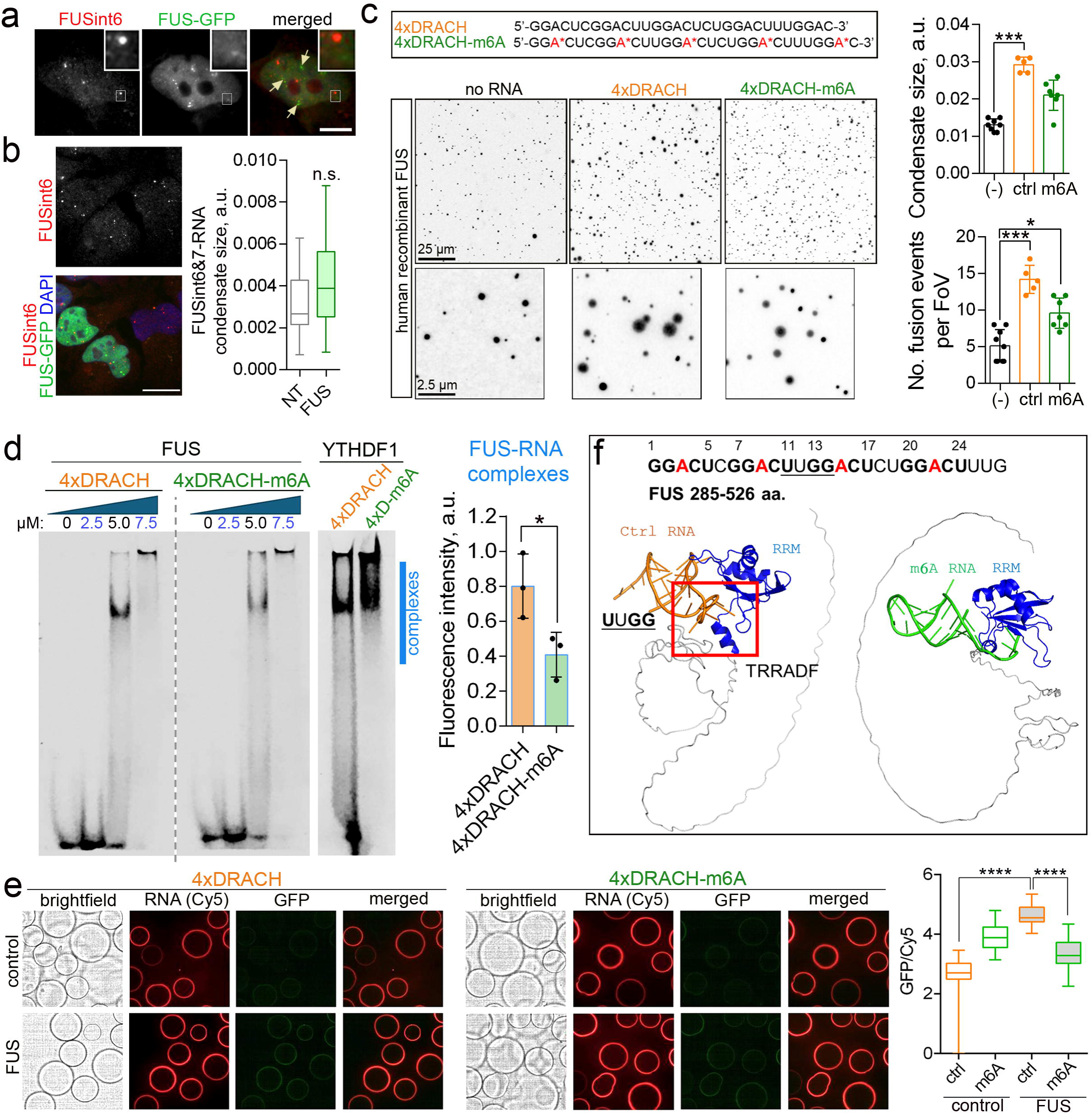
m6A has a repelling effect on FUS protein. **a** Ectopically expressed GFP-tagged FUS is not enriched in the endogenous FUSint6&7-RNA condensates. FUS intron 6 Stellaris probe was used in HeLa cells. Representative image is shown. Note FUS accumulation in paraspeckles (arrows). Scale bar, 5 µm. **b** FUS overexpression does not affect the FUSint6&7-RNA condensate size. Representative images and quantification for HeLa cells are shown. NT, non-transfected (cells analysed in the same FoV). Condensate size was measured in 15 FoV per condition. n.s., non-significant. Scale bar, 10 µm. **c** m6A limits FUS condensation *in vitro*. Recombinant human his-tagged FUS and synthetic Cy5-labelled RNA oligonucleotides, fully methylated or unmodified, were used. Representative images of FUS condensates and quantification of average condensate size and fusion events are shown. 5-8 FoV were analysed per condition from a representative experiment. *p<0.05, ***p<0.001, Kruskal-Wallis with Dunn’s test. **d** FUS-RNA complex formation is diminished by methylation, as analysed by EMSA. Recombinant FUS protein and RNA oligonucleotides as in *c* were used. Recombinant YTHDF1 protein was used in parallel as a positive control. Complex formation in the indicated area (blue line) was quantified by densitometry (5.0 and 7.5 μM data-points combined). N=3, *p<0.05, Mann-Whitney *U* test. **e** Confocal nanoscanning assay (CONA) demonstrates reduced binding of FUS to methylated RNA, as compared to its unmodified counterpart. Cy5-labelled RNA oligonucleotides as above and lysates of cells expressing GFP-tagged FUS were used. Representative images of beads and quantification of GFP ring fluorescence intensity (normalised by Cy5 fluorescence – RNA coating) are shown. 80-100 beads were analysed per condition using an automated pipeline. ****p<0.0001, Kruskal-Wallis with Dunn’s test. **f** AlphaFold3 modelling confirms the repelling properties of m6A in FUS binding to RNA. FUS RRM is in blue and RNA is in orange (unmodified) or green (methylated). Interacting interfaces of FUS protein and the DRACH motif-containing RNA are boxed. Amino acid sequence interacting with the UUGG motif in the oligonucleotide is also indicated.

We first utilised the *in vitro* ImmuCon assay for FUS condensate reconstitution that we recently developed (Hodgson et al., 2024), to test the effect of unmodified and m6A-modified RNA on FUS condensation. An RNA oligonucleotide containing 4 repeats of a DRACH motif GGACU (“4xDRACH”) and its fully modified counterpart (“4xDRACH-m6A”) were used (Fig. 7c). Recombinant FUS protein was mixed with RNA at a final concentration of 2.5 μM (previously found optimal for condensate formation, Hodgson et al., 2024), sedimented on glass coverslips, fixed and analysed by immunostaining. 4xDRACH RNA promoted FUS condensate formation, leading to large, frequently fusing condensates – suggestive of their high fluidity (Fig. 7c). Although 4xDRACH-m6A RNA did increase FUS condensation compared to the “no-RNA” control, the resultant condensates were significantly smaller and fused less as compared to the non-methylated RNA (Fig. 7c). m6A was previously reported to enhance phase-separation properties of RNA (Ries et al., 2019). Indeed, 4xDRACH-m6A underwent more efficient condensation than its non-modified counterpart (Supplementary Figure 7a), indicating that its homotypic condensation outcompeted co-condensation with FUS. Electrophoretic mobility shift assay (EMSA) also revealed more efficient formation of FUS-RNA complexes with 4xDRACH RNA as compared 4xDRACH-m6A RNA (Fig. 7d).

To further corroborate these findings, we used a version of the on-bead confocal nanoscanning assay (CONA) (Hintersteiner et al., 2010), where ring formation by a GFP-tagged RBP from cell lysate on RNA-coated beads can be used as a measure of its LLPS (Huang et al., 2024). We first confirmed that a positive control, m6A reader YTHDF1, readily accumulated on the beads coated with either unmodified or m6A-modified 4xDRACH RNA, with a clear preference for the latter (Supplementary Figure 7b). Analysis of FUS protein samples demonstrated diminished ring signal intensity on the beads coated with 4xDRACH-m6A RNA as compared to unmodified RNA (Fig. 7e).

Finally, we used AlphaFold3 to predict interactions between FUS RNA-binding sequences (C-terminus, aa.285-526) and the above RNA oligonucleotides. This analysis revealed an interaction between a sequence in the FUS RRM (aa. TRRADF) and the middle portion of 4xDRACH RNA (UUGG) but not its methylated counterpart (Fig. 7f).

These data suggest that FUS protein is excluded during condensation of m6A-modified RNA, reducing its enrichment in FUSint6&7-RNA condensates, possibly due to a direct repulsion by the m6A mark.

## Discussion

Our work argues that RNA condensation and delayed post-transcriptional splicing play a profound role in FUS expression. This regulation relies on a relatively abundant and stable FUS splicing intermediate that retains two conserved introns, 6 and 7. *First*, we demonstrate that this RNA species can assemble into higher-order ribonucleoprotein structures that contribute to FUS mRNA production. *Second*, our data suggest that gain-of-function mutations in FUS trigger a switch from a negative- to a positive-feedback autoregulation mode, with major implications for ALS pathology. *Finally*, we provide evidence that the retained introns themselves can be exploited to control FUS expression, opening up new therapeutic possibilities (Fig. 8).

**Figure 8.**
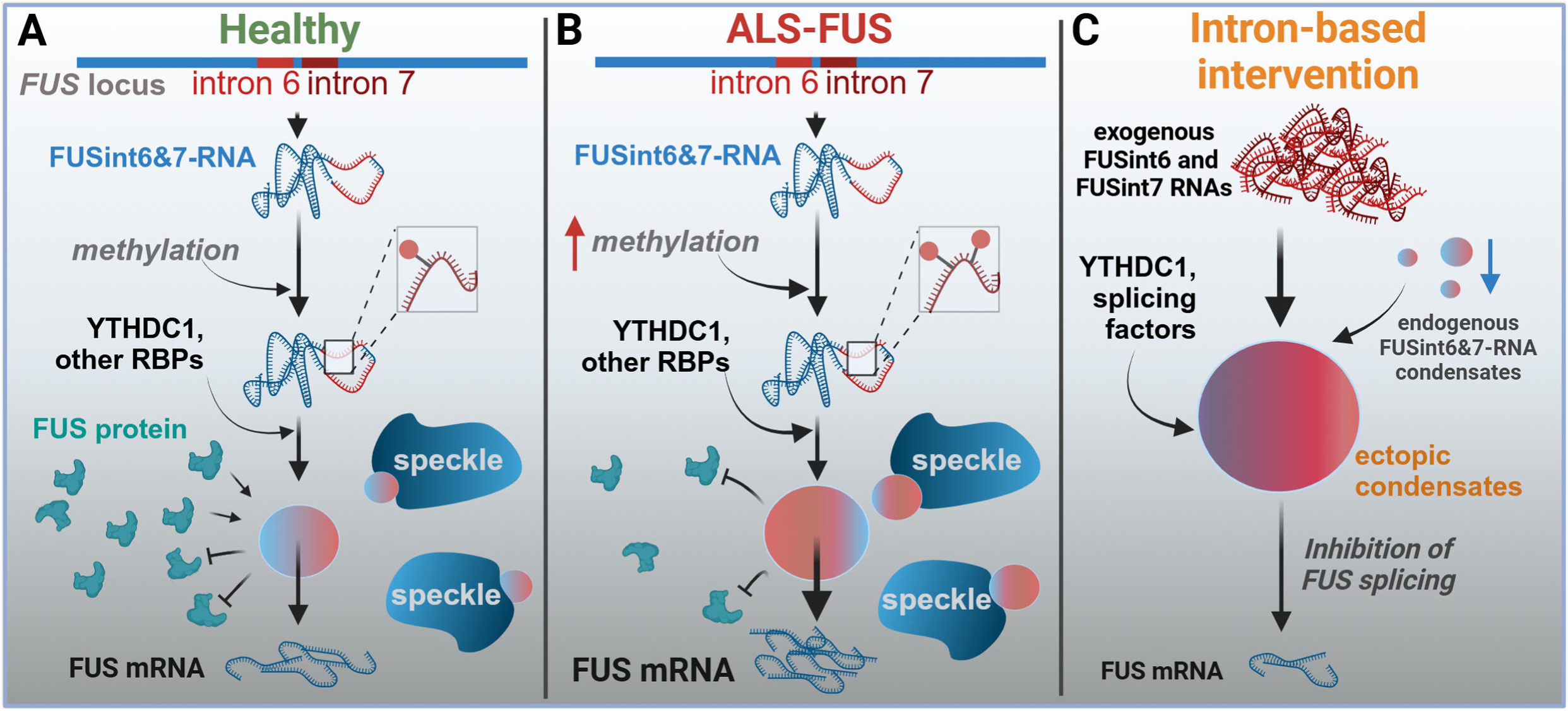
Regulation of FUS RNA condensation and post-transcriptional splicing in WT and ALS-FUS cells and the effect on exogenous FUS introns.

### Relationship between retained introns, RNA condensates, and FUS expression

We show that FUS transcripts retaining introns 6 and 7 form nuclear condensates in a variety of cell types, including neurons. In particular, FUS intron 7 in isolation is prone to forming microscopically visible foci and may thus drive the condensation behaviour of the entire transcript. RNA folding predictions indicated that intron 7 is more “structured” than non-foci-forming intron 6, featuring higher base-pairing probabilities and lower ensemble diversity. Indeed, highly structured RNAs were previously shown to form globular condensates, in contrast to “disordered” RNAs that tend to form mesh-like networks (Ma et al., 2021).

The characteristic number of large FUSint6&7-RNA foci per nucleus points to their assembly near the *FUS* transcription sites (Fig. 2). Yet, several lines of evidence suggest that these structures are distinct from the clusters of pre-mRNA molecules occasionally observed near transcription sites of actively expressed genes (e.g., *ACTB*; Supplementary Figure 5). *First,* these FUSint6&7-RNA foci are substantially larger than pre-mRNA clusters detected at the *ACTB* transcription sites (compare Fig. 2 and Supplementary Figure 5), despite the steady-state expression of *ACTB* is ∼18-fold higher than that of *FUS* (6761 nTPM *vs.* 376 nTPM, respectively; Human Protein Atlas). *Second*, smaller FUSint6&7-RNA condensates are often detected in addition to the main foci – also associated with speckles (Fig. 2, Fig. 3), whereas no such “secondary” condensates form in the case of *ACTB*. *Third*, FUSint6&7-RNA appears to be more abundant than fully unspliced FUS pre-mRNA species (Supplementary Figure 1), being sufficiently stable to act as a structural component of ribonucleoprotein condensates. The behaviour of FUSint6&7-RNA is reminiscent of that of NEAT1_2, a structural scaffold for paraspeckles. Paraspeckle spheroids cluster at the site of transcription, with a fraction of individual particles “budding” off this site and attaching to (but not fusing with) nuclear speckles (Takakuwa et al., 2023).

The proximity of FUSint6&7-RNA condensates to nuclear speckles (Fig. 3) and the abundance of splicing factors in their interactome/proteome (Fig. 5) supports a model in which these transcripts act as “reservoirs” of relatively long-lived splicing intermediates that can rapidly yield mature FUS mRNA, in a condition-dependent manner. Notably, post-transcriptional splicing accounts for ∼20% of splicing in human cells, often occurring near nuclear speckles (Girard et al., 2012; Wu et al., 2024). Dispersal of speckles by knocking down their core protein SON increases retention of speckle-associated introns (Barutcu et al., 2022). Thus, nuclear speckle proximity of the FUSint6&7-RNA condensates might facilitate the final steps of FUS mRNA biogenesis. Although further work will clearly be needed to assess the dynamic range of this “reservoir” mechanism, we propose that it may buffer gene expression changes under stress conditions. A recent study employing direct, full-length RNA sequencing in arsenite-treated cells has demonstrated that stress induces transcriptome-wide mRNA decay (Dar et al., 2024). Enhanced FUSint6&7-RNA splicing might therefore be important to maintain the FUS mRNA level for optimal recovery from stress.

Stimulus-dependent splicing of retained introns has been previously documented, including in neuronal contexts (Mauger et al., 2016; Mazille et al., 2022; Sung et al., 2023). Extending the scope of such regulation, a group of retained introns in mammals, termed “detained introns”, has been identified based on their ability to accumulate in the nucleus before undergoing post-transcriptional splicing (Boutz et al., 2015). The high interspecies conservation of FUS introns 6 and 7, their responsiveness to external stimuli, relative stability, nuclear retention, and NMD resistance, indicate that these introns belong to the above category. Interestingly, in line with detained introns responding to DNA damage (Boutz et al., 2015), HyPro-MS highlighted the presence of DNA damage/dsDNA break repair factors in FUSint6&7-RNA condensates.

We previously described the formation of phase-separated structures in the nucleus for Srsf7 RNA with retained introns (Königs et al., 2020). More recently, we have systematically catalogued condensate-forming RNAs and identified hundreds of RNAs with conserved retained introns (Hernandez-Canas, Busam et al., *manuscript in preparation).* Future studies should explore the extent to which condensation of stable RNA species with retained introns contributes to post-transcriptional splicing regulation and the dynamics of nuclear RNA “reservoirs”.

### ALS-FUS mutations may rewire FUS RNA condensate regulation

An important finding of our study is the role of the m6A reader YTHDC1 in the FUSint6&7-RNA condensate regulation (Fig. 6). m6A modification is emerging as a key factor in RNA compartmentalisation and fate in the nucleus. For example, m6A modification of MALAT1 significantly modulates speckle composition and properties (Wang et al., 2021), whereas enhancer RNA methylation facilitates enhancer RNA (eRNA) condensate assembly (Lee et al., 2021). m6A mark has a direct condensation-promoting effect *in vitro* (Ries et al., 2019). In cells, YTHDC1, which has two intrinsically disordered regions and is capable of phase separation, likely also contributes to RNA condensation (Lee et al., 2021; Widagdo et al., 2022). Furthermore, RNA condensation via m6A/YTHDC1 was reported to promote nuclear retention of RNA (Lee et al., 2025) and protect RNA from degradation (Cheng et al., 2021). Our study expands the repertoire of m6A/YTHDC1-dependent nuclear condensates.

Previous studies proposed two negative-feedback mechanisms maintaining FUS homeostasis (Humphrey et al., 2020; Zhou et al., 2013). These mechanisms rely on FUS protein binding to the intron 6 - exon 7 - intron 7 region of its pre-mRNA, promoting either exon skipping or intron retention. While loss of wild-type FUS in our KO cell model did not alter FUSint6&7-RNA abundance (Fig. 1), our intron 7 overexpression studies suggested that FUS can still be recruited onto its own transcript (Fig. 5), consistent with the published iCLIP data (Rogelj et al., 2012). Intriguingly, unlike FUS KO, ALS-FUS mutations had a suppressive effect on FUSint6&7-RNA in our studies (Fig. 1). This aligns with reduced intron 6/7 retention observed in three ALS-FUS mouse models, but not in three FUS KO models in a previous study (Fig. 4c in Humphrey et al., 2020). Our data on m6A modification may explain this behaviour. Indeed, FUS RNA methylation levels and YTHDC1 recruitment significantly increase in ALS-FUS mutant cells (Fig. 6). This in turn potentiates the assembly of FUSint6&7-RNA condensates – positive regulators of FUS post-transcriptional splicing (Fig. 6). Simultaneously, m6A-mediated condensation may hinder FUS protein access to its binding sites within the intron 6 - exon 7 - intron 7 region (Fig. 7). We propose that ALS-FUS mutations rewire the post-transcriptional regulation of FUS, disrupting its self-inhibitory feedback loop and at the same time inducing abnormal self-activation (Fig. 8a,b). FUS upregulation is typical for both ALS-FUS models and patient tissue (An et al., 2019; Lashley et al., 2011; Sabatelli et al., 2013), and even a modest increase in FUS level is toxic *in vivo* (Ling et al., 2019; Mitchell et al., 2013). Thus, the mutation-induced switch in the FUS regulatory logic will have profound implications for its expression control and ALS pathology.

An important question for future studies is why ALS-FUS mutations increase m6A modification of FUS RNA. One possibility is that the mutation-induced shift of FUS protein to the cytoplasm leaves its RNA more exposed to m6A writers. Alternatively, this effect may arise through indirect mechanisms, including disruption to RBP regulatory networks (Khoroshkin et al., 2024; Koike et al., 2023; Martinez et al., 2016; Ziff et al., 2023). Indeed, mutant FUS has been shown to reduce the solubility of multiple RBPs (Korobeynikov et al., 2022).

### Therapeutic potential of FUS introns

Recently, a study by the Dupuis group has described a striking amelioration of mutant FUS toxicity in mice by introduction of a full human *FUS* transgene, including the rescue of lethality/shortened lifespan and paralysis in homozygous and heterozygous Fus^ΔNLS^ mice, respectively (Sanjuan-Ruiz et al., 2021). This was associated with accumulation of endogenous FUSint6&7-RNA and reduced (mutant) FUS mRNA and protein production. Phenotype rescue in these double-transgenic mice seemed counter-intuitive, since FUS overexpression was toxic in other FUS transgenic mice (Ling et al., 2019; Mitchell et al., 2013). The key difference in the study design was the presence of all *FUS* regulatory sequences in the transgene used in this recent study.

Our data (Fig. 4) argue that FUS intron 7 may be responsible for the therapeutic effect of the full *FUS* transgene. Indeed, exogenous expression of FUS intron 7 – alone or in combination with intron 6 – leads to ectopic condensate formation and sequestration of certain splicing factors, YTHDC1, and potentially, FUSint6&7-RNA itself (Fig. 8c). Depletion of these factors results in the dissolution of endogenous condensates, inhibited post-transcriptional splicing and ultimately, FUS mRNA and protein downregulation. Coincidentally, FUS ASO Jacifusen targets FUS intron 6 (Korobeynikov et al., 2022), therefore both FUS pre-mRNA and FUSint6&7-RNA are expected to be degraded. Thus, FUSint6&7-RNA (condensate) depletion may contribute to efficient FUS mRNA downregulation by this ASO. Strikingly, in our cellular studies, even high levels of exoFUSint6/7-RNA were not cytotoxic. It is possible that these ectopic condensates sequester a limited subset of splicing factors, causing no major changes to the global splicing patterns. Future studies should verify the above phenotypes in neuronal models using viral delivery vectors. FUS introns are relatively long, and it would be important to establish the minimal regulatory sequence(s).

Our studies pinpoint a central regulatory element in the *FUS* gene that can confer a therapeutic effect and opens a novel avenue for FUS modulation in ALS-FUS. The homeostatic nature of this mechanism, targeting an endogenous regulatory switch, and the ability to establish permanent expression using a gene therapy vector are the obvious advantages. Furthermore, it may be possible to target FUSint6&7-RNA condensates using specific small molecules (“c-mods”) (Patel et al., 2022). Such approaches would offer a novel therapeutic strategy leveraging the mechanisms we have described.

## Materials and Methods

### Splice site analysis

Sequence of the human *FUS* gene (transcript ENST00000254108.12/NM_004960.4) was downloaded from Ensembl (Harrison et al., 2024) with introns and exons annotated. For each splice acceptor and splice donor site, the input sequence required to run MaxEntScan (Yeo and Burge, 2004) (acceptor: 20 bases intronic sequence, 3 bases exonic, donor: 3 bases exonic, 6 bases intronic sequence) was manually extracted and stored in fasta format. The acceptor and donor fasta files were run through the relevant MaxEntScan webservers (acceptor: http://hollywood.mit.edu/burgelab/maxent/Xmaxentscan_scoreseq_acc_fugu.html, donor: http://hollywood.mit.edu/burgelab/maxent/Xmaxentscan_scoreseq_fugu.html) with scoring using the Maximum Entropy Model.

### FUS RNA isoform analysis

The Sequence Read Archive (SRA) was used to obtain RNA-Seq from human cerebellar tissue from an unaffected individual (project accession: PRJNA279249, run accession: SRR1927061). Reads were aligned to GRCh38 with annotations from GENCODE v47 using STAR run via Galaxy (usegalaxy.org, version 24.1.4.dev0), with default settings. Aligned sequencing reads were visually inspected using the Integrative Genomics Viewer (IGV).

### Expression constructs

Plasmids for ectopic expression of FUS introns were generated by amplifying the intron sequences with a portion of the flanking exon sequences, preserving the splice site; primers are given in Supplementary Table S4. Resultant fragments were inserted in pEGFP-N1 vector between XhoI and KpnI sites (upstream of GFP ORF). Plasmids for GFP-tagged WT and mutant FUS expression were described previously (Shelkovnikova et al., 2014a; Shelkovnikova et al., 2014b).

### Cell culture experiments

SH-SY5Y, HeLa and U2OS WT cells from ECACC repository were maintained in DMEM/F12 media supplemented with 10% foetal bovine serum and 1x Penicillin-Streptomycin-Glutamine (all from Invitrogen). FUSΔNLS lines were generated and characterised previously (An et al., 2019). Cells were plated on uncoated 10-mm coverslips in 24-well plates and left to recover overnight before transfection or treatments. Cells were transfected with plasmid DNA using either Lipofectamine2000 (Invitrogen) or JetPRIME reagent (Sartorius), according to manufacturer’s instructions. siRNA transfections were done using Lipofectamine2000. Silencer®Select validated siRNAs for YTHDC1 and METTL16 were purchased from Thermo Scientific. AllStars scrambled siRNA (Qiagen) was used as a control. The following chemical modulators of cellular pathways were used: sodium arsenite (0.5 mM), cycloheximide (10 μM), actinomycin D (2.5 μg/ml), interferon-beta (all Sigma); pladienolide-B (200 nM), STM2457 (10 μM), and FB23-2 (10 μM) (all Cayman Chemicals). Cell survival analysis was performed 24 h post-transfection using CellTiter-Blue® Cell Viability Assay (Promega) according to manufacturer’s instruction, in a 96-well format, after verification of transfection efficiency by imaging. For 1,6-hexanediol, DNase I and RNase A treatments in semi-permeabilised cells, HeLa cells were treated with 1% Tween/PBS for 10 min before subjecting to a treatment: 5% 1,6-hexanediol in PBS for 5 min, 0.2 mg/mL DNase I or RNase A (Qiagen) diluted in PBS, for 20 min, followed by fixation as normal. Human motor neurons were differentiated from human ES cells (H9) and maintained as described in Shelkovnikova et al., 2017.

### Generation of cell clones with FUS intron 6 partial deletion

For generation of cell clones with FUS intron deletion (Δint6), sgRNA sequences were selected using the IDT gRNA online tool: forward, AACCATTACCAACTCCCTAG and reverse, TGTCAACCACTTGCGACAAG. Annealed DNA oligonucleotides (custom-made by Merck) were cloned into pX330 vector (Cong et al., 2013) (Addgene plasmid #42230). HeLa cells were transfected with both plasmids using jetPRIME, subjected to limited dilution, and single cell-derived clones were analysed by PCR (primers are given in Supplementary Table S4) using Go-Taq Green Master Mix (Promega). Genomic DNA was isolated by proteinase K (Sigma) digest of cell pellets. Successful editing was confirmed by DNA sequencing.

### RNA fluorescence in situ hybridisation (RNA-FISH) and immunocytochemistry

Cell fixation, processing, RNA-FISH and immunocytochemistry on coverslips were performed as described earlier (Shelkovnikova et al., 2018). Fluorescently labelled RNA-FISH probes for FUS introns 6, 7 and 1 were designed and custom-made by Biosearch Technologies (Stellaris® probes). Oligonucleotide sequences provided in Supplementary Table S5. FUS intron 6 and 7 probes were labelled with Quasar®570, and intron 1 probe was labelled with FITC. The single DNA oligonucleotide probe used for exoFUSint7 condensate detection was 5’-labelled with Cy5: 5’-TCCCGAGGGCCTTTAGTGAC-3’ (custom-made by Merck). A 5’-Cy5-labelled DNA oligonucleotide was also used for FUS intron 1 detection: 5’-CGCTTCAAACCCCTAAGAAG-3’. Paraspeckles were detected using a NEAT1_2 specific Stellaris® custom probe labelled with Quasar®670. RNA-FISH in a 96-well format was carried out as described before (An et al., 2022), except the final probe concentration was 50 nM. For ICC, cells prepared as for RNA-FISH were washed with 1xPBS, incubated in primary antibody (1:1000) for 2 h and subsequently in secondary antibody (AlexaFluor, Invitrogen, 1:1000) for 1 h at RT. The following commercial primary antibodies were used: FUS (mouse monoclonal, Santa Cruz, sc-47711 and rabbit polyclonal, Proteintech, 11570-1-AP); SMN (mouse monoclonal, BD Biosciences, 610646); coilin p80 (mouse monoclonal, BD Biosciences, 612074); TDP-43 (rabbit polyclonal, C-terminal, Sigma); YTHDC1 (rabbit polyclonal, Proteintech, 14392-1-AP); HDGFL2 (rabbit polyclonal, Proteintech, 15134-1-AP); hnRNPC (rabbit polyclonal, Proteintech, 11760-1-AP); PNN/pinin (rabbit polyclonal, Proteintech, 18266-1-AP); ANP32B (rabbit polyclonal, Proteintech, 10843-1-AP); FUBP1 (rabbit polyclonal, Proteintech, 24864-1-AP). For imaging on coverslips, Olympus BX57 fluorescent microscope (x100 oil objective) equipped with ORCA-Flash 4.0 camera (Hamamatsu) and cellSens Dimension software (Olympus) was used. Imaging on Opera Phenix was performed at 80% laser power and 500 ms exposure for Cy3 channel. FUS RNA condensate properties (area, number, solidity, fluorescence intensity) were quantified using Analyze particles tool in Image J https://imagej.net/ij/. Large condensates were determined as foci >50px or >100px in size, depending on the detection efficiency in a given experiment.

### RNAscope®-ISH analysis

Cells cultured on coverslips were fixed in 10% neutral buffered formalin (NBF), dehydrated through ascending ethanol concentrations and kept in 100% ethanol at −20⁰C until use. On the day of assay, coverslips were rehydrated, washed with PBS, subjected to protease IV (ACDBio) pre-treatment for 15 min at RT, washed again and incubated with a custom FUS intron 6 RNAscope probe for 2 h at 40⁰C. The probe was designed and manufactured by ACDBio/Bio-Techne (design #NPR-0039133). Signal was detected using RNAscope™ 2.5 HD Assay - BROWN kit according to manufacturer’s instructions. Coverslips were mounted for imaging using Immu-Mount (ThermoFisher). Images were taken using either Olympus BX57 with ORCA-Flash 4.0 camera or Nikon Eclipse Ni equipped with Nikon DS-Ri1 camera.

### HyPro-MS analysis of FUS RNA condensate composition

Digoxigenin-labelled DNA probe sets specific for FUSint6&7 were designed using Stellaris online tool and custom-made by Eurofins (Supplementary Table S5). Probes were labelled using a 2nd generation DIG Oligonucleotide 3′ End Labeling Kit (Sigma Aldrich) to yield 5 μM digoxigenin-labeled mixtures. Recombinant HyPro enzyme was prepared and HyPro labeling was performed as described (Yap et al., 2022b). Briefly, cells grown in 10-cm dishes or on coverslips were fixed with 0.5 mg/ml dithiobis(succinimidyl propionate) (DSP; Thermo Fisher Scientific) in 1xPBS for 30 min at RT. The samples were then washed with 1xPBS / 20 mM Tris-HCl, pH 8.0, permeabilised with 70% ethanol, equilibrated in 2xSSC and 10% formamide (ThermoFisher Scientific), and hybridized with digoxigenin-labeled probes (100 nM for FUSint6&7 and 150 nm for ACTB) in 2xSSC, 10% formamide and 10% dextran sulfate overnight at 37°C. Samples were washed with 10% formamide in 2xSSC and at 37°C for 30 min and 1xSSC at RT for 15 min and blocked with 0.8% BSA in 4xSSC (HyPro blocking buffer) treated with 100 murine RNase inhibitor (New England Biolabs). “Samples were incubated with 2.7 μg/ml of enhanced HyPro enzyme in HyPro blocking buffer at RT for 1 h. After washing off unbound HyPro, proximity biotinylation was carried out in the presence of 0.5 mM biotin-phenol (Caltag Medsystems) and 0.1 mM hydrogen peroxide (Sigma Aldrich) as described (*Yap et al., submitted*). The reaction was quenched by 5 mM Trolox (Sigma Aldrich) and 10 mM sodium ascorbate (Sigma Aldrich, cat# A4034) in 1xPBS. Samples labelled in dishes were then analysed by immunoblotting, mass-spectrometry or qRT-PCR. The coverslips were used for HyPro-FISH. Cells on dishes were lysed in high-SDS RIPA buffer with 10 mM sodium ascorbate, 5 mM Trolox, 50 mM DTT, cOmplete, EDTA-free protease inhibitor cocktail (Sigma Aldrich), and 1 mM PMSF (Cell Signaling), sonicated, de-crosslinked and cleared by centrifugation. MyOne streptavidin C1 magnetic beads were incubated with de-crosslinked lysates for 1 h at RT. The beads were washed twice with RIPA buffer, once with 1 M KCl, once with 0.1 M Na2CO3, once with 2 M urea in 10 mM Tris-HCl, pH 8.0 and twice with RIPA buffer. The beads were collected using DynaMag-2 Magnet and analysed by mass spectrometry performed by the CEMS Proteomics Core Facilities at King’s College London as described (Yap et al., 2022b). Raw mass-spec data files were processed using Proteome Discoverer (v2.2; Thermo Fisher Scientific) to search against Uniprot Swissprot Homo sapiens Taxonomy (49,974 entries) using Mascot (v2.6.0; www.matrixscience.com) and the Sequest search algorithms. Precursor mass tolerance was set to 20 ppm with fragment mass tolerance set to 0.8 Da with a maximum of two missed cleavages. Variable modifications included carbamidomethylation (Cys) and oxidation (Met). Searching stringency was set to 1% False Discovery Rate (FDR). In total, 2869 proteins were detected. LFQ intensity output was filtered against proteins commonly found in proteomic contamination database CRAPome (Mellacheruvu et al., 2013). Keratins were also removed. The filtered data was then then imported to R and analysed using the DEP package (https://bioconductor.org/packages/release/bioc/vignettes/DEP/inst/doc/DEP.html). The data were filtered to include the proteins only identified in all 3 replicates, with default imputation settings (fun = “MinProb”, q = 0.01). DEP-generated P-values were adjusted for multiple testing using the Benjamini-Hochberg (FDR) method.

### RNA purification and qRT-PCR

RNA was extracted either from total cell lysates or from nuclear and cytoplasmic fractions using TRI-reagent (Sigma) with a heating step (55⁰C for 10 min). REAP protocol was used for nuclear-cytoplasmic fractionation (Suzuki et al., 2010). In semi-extractability analysis, non-heated samples (kept at RT) were analysed in parallel (An et al., 2019). First-strand cDNA synthesis was performed using 500 ng of RNA with random primers (Promega) and MMLV reverse transcriptase (Promega) as per manufacturer’s protocol (25 µl final reaction volume). qRT-PCR was run using qPCRBIO SyGreen Lo-ROX mix on CFX96/C1000™ qPCR system (Bio-Rad). Samples were analysed in at least 2 technical repeats, and expression of specific genes was determined using the 2^−ΔΔCt^ method and GAPDH for normalisation. Primer sequences are given in Supplementary Table S4. Primers were designed using the NCBI Primer design tool (BLAST+Primer3).

### Western blotting

Total cell lysates were prepared by adding 2x Laemmli buffer directly to the wells in a 24-well plate followed by denaturation at 95°C for 10 min. SDS-PAGE and detection of proteins were carried out as described before (Huang et al., 2024). The following commercial primary antibodies were used (1:1000 dilution): FUS (rabbit polyclonal, 11570-1-AP, Proteintech and rabbit polyclonal, Bethyl, A300-293); YTHDC1 (rabbit polyclonal, Proteintech, 29441-1-AP).

### Analysis of global m6A RNA levels

For dot blot, 500 ng of total RNA was UV-crosslinked to Hybond-N+ membrane (Merck), which was subsequently probed with an m6A-specific antibody (mouse monoclonal, Proteintech, 68055-1-Ig) as described for western blot. Results were validated using an independent antibody (rabbit polyclonal, Cell Signaling, 56593). EpiQuik m6A RNA Methylation Quantification Kit (EpigeneTek, P-9005) was used on the same set of RNA samples (200 ng of RNA per well) with inclusion of additional controls (FUS KO and FUS-GFP expressing cells), according to the manufacturer’s instructions.

### RNA immunoprecipitation (RIP)

WT and FUSΔNLS SH-SY5Y cells were transfected with pcDNA3-FLAG-HA-hYTHDC1 plasmid (gift from Samie Jaffrey, Addgene plasmid #85167) (Patil et al., 2016) or control plasmid (Venus-Flag, made in-house) and harvested 24 h post-transfection in the IP buffer (1xPBS, 1% Triton-X100 with cOmplete Mini Protease Inhibitor Cocktail). Transfection efficiency in all cell lines was verified by western blot. After 30 min lysis on ice with periodic vortexing, the lysates were cleared by centrifuging at 13,000rpm for 20 min. ChromoTek DYKDDDDK Fab-Trap® Agarose beads were added to each supernatant sample and incubated for 3 h on nutator at 4⁰C. Beads were washed with IP buffer three times, and bound RNAs were eluted in TRI-reagent. RNA purification, cDNA synthesis and qRT-PCR were performed as above, using 150 ng RNA as a template. RNA levels were normalised to GAPDH mRNA.

### Methylated RNA immunoprecipitation (MeRIP)

Total RNA was isolated from WT and FUSΔNLS SH-SY5Y cells using TRI-reagent with heating. For immunoprecipitation, 6 μg of a mouse anti-m6A antibody (mouse monoclonal, Proteintech, 68055-1-Ig) was incubated with 25 μl Dynabeads Protein G (Invitrogen, 10003D) in IP buffer (10mM Tris pH 7.5, 150mM NaCl, 0.1% NP-40) for 1 h at 4⁰C, followed by washes with the same buffer. Subsequently, 30 μg of total RNA was added to the m6A antibody coated beads for 2 h at 4⁰C. Uncoated beads were used as a control. After IP buffer washes, the beads were resuspended in TRI-reagent, followed by RNA isolation. cDNA synthesis and qPCR were performed as above, using 170 ng RNA as a template. RNA levels were normalised to GAPDH mRNA and subsequently to the no-antibody control.

### Analysis of FUS condensation in vitro by ImmuCon

ImmuCon Assay is described in detail in Hodgson et al., 2024. Recombinant FUS (produced in-house; 2.5 μM final concentration) was mixed with 250 nM unmodified or m6A-modified 4XDRACH RNA oligonucleotide in assay buffer (20mM Tris-HCl pH7.5, 20 mM KCl, 2 mM MgCl2, 1 mM DTT). RNA oliogonucleotides (5’-end labelled with Cy5 and 3’-end labelled with biotin-TEG) were custom-made and HPLC-purified by Horizon (*denotes m6A-modified adenosine): Ctrl, 5’-GGACUCGGACUUGGACUCUGGACUUUGGAC-3’ and m6A, 5’-GGA*CUCGGA*CUUGGA*CUCUGGA*CUUUGGA*C-3’. A microdroplet (5 μl) of each sample was placed on a 10-mm round glass coverslip; condensates were left to sediment for 15 min and then fixed with 2% glutaraldehyde in PBS for 15 min. The coverslips were blocked by 1% BSA in PBS for 1 h at RT and subsequently incubated with an anti-FUS antibody (1:5000, rabbit polyclonal, Proteintech 11570-1-AP) for 2 h at RT. After PBS washes, an anti-rabbit AlexaFluor546 antibody (1:1000, Invitrogen) was applied for 1 h. The coverslips were mounted with Immu-mount (ThermoFisher) and imaged using Olympus BX57 upright microscope and ORCA-Flash 4.0 camera and processed using cellSens Dimension software (Olympus). FUS condensate area was quantified using the Analyze Particles tool in ImageJ.

### Confocal nanoscanning (CONA)

CONA assay was performed as described for TDP-43 (Huang et al., 2024), with modifications. Plasmid for YTHDF1-GFP expression was a gift from Elisa Izaurralde (pT7-EGFP-C1-HsYTHDF1_AB, Addgene plasmid #148307). Briefly, HeLa cells transfected to express either GFP alone or GFP-tagged FUS or YTHDF1 in 35-mm dishes were lysed in 500 µl of 1% Triton-X100/PBS with 40U/ml RiboLock (ThermoFisher) for 15 min on ice with periodic vortexing. Lysates were cleared by centrifuging at 13,000 rpm for 15 min, snap-frozen and kept in −80°C. Ni-NTA Superflow beads (Qiagen) were washed with binding buffer (20 mM HEPES pH 7.5, 0.3 M NaCl, 0.01% Triton X-100, 5 mM MgCl_2_), coated with His-streptavidin (NKMAX) and then with the respective RNA oligonucleotide described above – 4xDRACH or 4xDRACH-m6A (40 pmol), or “generic” RNA oligonucleotide mix (a mix of (UG)_15_, (AUG)_12_, Clip34nt, in equal ratios). Beads were washed in binding buffer and added to the thawed cell lysates containing the GFP-tagged protein and incubated on multi-orbital rotator for 2 h at RT. After washes in binding buffer, beads were imaged in µCLEAR 384 well plate (Greiner) on Opera Phenix. Mean ring intensity for Cy5 and EGFP channels was quantified using a custom pipeline on Harmony 4.9.

### Electrophoretic mobility shift assay (EMSA)

Cy5-labelled RNA oligonucleotides 4xDRACH and 4xDRACH-m6A were incubated at 250 nM with recombinant FUS (as above) at 0, 2.5, 5.0 and 7.0 μM or YTHDF1 (Active Motif #31608; at 0.64 μM) in the assay buffer (50 mM Tris-HCl, pH 7.5, 100 mM KCl, 2 mM MgCl_2_, 100 mM β-mercaptoethanol, 0.1 mg/ml BSA) for 30 min at RT with gentle shaking. Samples were analysed on 6% native acrylamide gel in TBE buffer, followed by imaging on Licor Odyssey FC (700nm channel). Band intensities were quantified by ImageStudio.

### Analysis of FUS RNA methylation

Datasets for the human *FUS* gene publicly available as part of the m6A-Atlas 2.0 http://rnamd.org/m6a/ (Liang et al., 2024) were used for analysis. The full sequence of *FUS* from Ensemble database (gene ID ENST00000254108.12), viewed in the UCSC Genome Browser on human chr16:31,180,139-31,191,605 (GRCh38/hg38), was used for alignment.

### In silico analysis of FUS intron structure

For FUS introns 6 and 7 structure prediction and visualisation, RNAfold v2.0 (Lorenz et al., 2011) was used, with the parameters described earlier (Mathews et al., 2004).

### FUS – RNA interaction predictions

Predictions of FUS interaction with unmodified and m6A-modified 4xDRACH oligonucleotides were performed using AlphaFold3. The top-ranked results were output and visualised in Pymol v.2.55.5.

### Data visualisation and statistics

Processed data visualisation and statistical analysis were performed using GraphPad Prism software unless indicated otherwise. Mean values of replicates were compared using appropriate statistical tests. Statistical tests used are indicated in figure legends with statistical significance denoted with asterisks: *p<0.05, **p<0.01, ***p<0.001, ****p<0.0001. N indicates the number of biological replicates. Error bars represent standard deviation (SD) unless indicated otherwise.

## Acknowledgements

The work was supported by the UKRI Future Leaders Fellowship (MR/W004615/1), MRC (MR/W028522/1) and BBSRC (BB/V014110/1) standard grants, and MND Association grant (968-799) to T.A.S.; and BBSRC grants BB/Y009304/1, BB/V006258/1, and BB/R001049/1 to E.V.M. E.D., B.S.C.E. and V.K. are supported by PhD studentships from MND Scotland, the University of Sheffield Faculty of Health and Neuroscience Institute, respectively (to T.A.S.).

## Author contributions

W.-P.H. designed and conducted experiments, analysed the data and contributed to the study supervision; V.K., K.Y., H.A., S.J.J., R.E.H., A.A.S., E.D., B.C.S.E., and T.H.C. conducted experiments and analysed the data. J.L. and M.M.-M. contributed to the study design and analysed the data; E.V.M. contributed to the study design, data analysis and study supervision and wrote the manuscript; T.A.S. conceived, designed and supervised the study, designed and conducted experiments, analysed data and wrote the manuscript. All authors contributed to the manuscript writing and approved its final version.

## References

An, H., Elvers, K.T., Gillespie, J.A., Jones, K., Atack, J.R., Grubisha, O., and Shelkovnikova, T.A. (2022). A toolkit for the identification of NEAT1_2/paraspeckle modulators. Nucleic Acids Res 50, e119.

An, H., Skelt, L., Notaro, A., Highley, J.R., Fox, A.H., La Bella, V., Buchman, V.L., and Shelkovnikova, T.A. (2019). ALS-linked FUS mutations confer loss and gain of function in the nucleus by promoting excessive formation of dysfunctional paraspeckles. Acta Neuropathol Commun 7, 7.

Barutcu, A.R., Wu, M., Braunschweig, U., Dyakov, B.J.A., Luo, Z., Turner, K.M., Durbic, T., Lin, Z.Y., Weatheritt, R.J., Maass, P.G., et al. (2022). Systematic mapping of nuclear domain-associated transcripts reveals speckles and lamina as hubs of functionally distinct retained introns. Molecular cell 82, 1035–1052 e1039.

Baumer, D., Hilton, D., Paine, S.M., Turner, M.R., Lowe, J., Talbot, K., and Ansorge, O. (2010). Juvenile ALS with basophilic inclusions is a FUS proteinopathy with FUS mutations. Neurology 75, 611–618.

Boutz, P.L., Bhutkar, A., and Sharp, P.A. (2015). Detained introns are a novel, widespread class of post-transcriptionally spliced introns. Genes Dev 29, 63–80.

Bresson, S., Shchepachev, V., Spanos, C., Turowski, T.W., Rappsilber, J., and Tollervey, D. (2020). Stress-Induced Translation Inhibition through Rapid Displacement of Scanning Initiation Factors. Molecular cell 80, 470–484 e478.

Cheng, Y., Xie, W., Pickering, B.F., Chu, K.L., Savino, A.M., Yang, X., Luo, H., Nguyen, D.T., Mo, S., Barin, E., et al. (2021). N(6)-Methyladenosine on mRNA facilitates a phase-separated nuclear body that suppresses myeloid leukemic differentiation. Cancer Cell 39, 958–972 e958.

Chujo, T., Yamazaki, T., and Hirose, T. (2016). Architectural RNAs (arcRNAs): A class of long noncoding RNAs that function as the scaffold of nuclear bodies. Biochim Biophys Acta 1859, 139–146.

Chujo, T., Yamazaki, T., Kawaguchi, T., Kurosaka, S., Takumi, T., Nakagawa, S., and Hirose, T. (2017). Unusual semi-extractability as a hallmark of nuclear body-associated architectural noncoding RNAs. The EMBO journal 36, 1447–1462.

Clark, M.B., Johnston, R.L., Inostroza-Ponta, M., Fox, A.H., Fortini, E., Moscato, P., Dinger, M.E., and Mattick, J.S. (2012). Genome-wide analysis of long noncoding RNA stability. Genome Res 22, 885–898.

Cong, L., Ran, F.A., Cox, D., Lin, S., Barretto, R., Habib, N., Hsu, P.D., Wu, X., Jiang, W., Marraffini, L.A., et al. (2013). Multiplex genome engineering using CRISPR/Cas systems. Science 339, 819–823.

Cook, K.B., Kazan, H., Zuberi, K., Morris, Q., and Hughes, T.R. (2011). RBPDB: a database of RNA-binding specificities. Nucleic Acids Res 39, D301–308.

Dani, C., Piechaczyk, M., Audigier, Y., El Sabouty, S., Cathala, G., Marty, L., Fort, P., Blanchard, J.M., and Jeanteur, P. (1984). Characterization of the transcription products of glyceraldehyde 3-phosphate-dehydrogenase gene in HeLa cells. Eur J Biochem 145, 299–304.

Dar, S.A., Malla, S., Martinek, V., Payea, M.J., Lee, C.T., Martin, J., Khandeshi, A.J., Martindale, J.L., Belair, C., and Maragkakis, M. (2024). Full-length direct RNA sequencing uncovers stress-granule dependent RNA decay upon cellular stress. eLife13:RP96284

Deng, H., Gao, K., and Jankovic, J. (2014). The role of FUS gene variants in neurodegenerative diseases. Nat Rev Neurol 10, 337–348.

Devoy, A., Kalmar, B., Stewart, M., Park, H., Burke, B., Noy, S.J., Redhead, Y., Humphrey, J., Lo, K., Jaeger, J., et al. (2017). Humanized mutant FUS drives progressive motor neuron degeneration without aggregation in ‘FUSDelta14’ knockin mice. Brain 140, 2797–2805.

Edupuganti, R.R., Geiger, S., Lindeboom, R.G.H., Shi, H., Hsu, P.J., Lu, Z., Wang, S.Y., Baltissen, M.P.A., Jansen, P., Rossa, M., et al. (2017). N(6)-methyladenosine (m(6)A) recruits and repels proteins to regulate mRNA homeostasis. Nat Struct Mol Biol 24, 870–878.

Girard, C., Will, C.L., Peng, J., Makarov, E.M., Kastner, B., Lemm, I., Urlaub, H., Hartmuth, K., and Luhrmann, R. (2012). Post-transcriptional spliceosomes are retained in nuclear speckles until splicing completion. Nat Commun 3, 994.

Grassano, M., Brodini, G., De Marco, G., Casale, F., Fuda, G., Salamone, P., Brunetti, M., Sbaiz, L., Gallone, S., Cugnasco, P., et al. (2022). Phenotype Analysis of Fused in Sarcoma Mutations in Amyotrophic Lateral Sclerosis. Neurol Genet 8, e200011.

Gruber, A.R., Lorenz, R., Bernhart, S.H., Neubock, R., and Hofacker, I.L. (2008). The Vienna RNA websuite. Nucleic Acids Res 36, W70–74.

Harrison, P.W., Amode, M.R., Austine-Orimoloye, O., Azov, A.G., Barba, M., Barnes, I., Becker, A., Bennett, R., Berry, A., Bhai, J., et al. (2024). Ensembl 2024. Nucleic Acids Res 52, D891–D899.

Hewitt, C., Kirby, J., Highley, J.R., Hartley, J.A., Hibberd, R., Hollinger, H.C., Williams, T.L., Ince, P.G., McDermott, C.J., and Shaw, P.J. (2010). Novel FUS/TLS mutations and pathology in familial and sporadic amyotrophic lateral sclerosis. Arch Neurol 67, 455–461.

Hintersteiner, M., Ambrus, G., Bednenko, J., Schmied, M., Knox, A.J., Meisner, N.C., Gstach, H., Seifert, J.M., Singer, E.L., Gerace, L., et al. (2010). Identification of a small molecule inhibitor of importin beta mediated nuclear import by confocal on-bead screening of tagged one-bead one-compound libraries. ACS Chem Biol 5, 967–979.

Hodgson R., Huang W.P., Kumar V., Haiyan A., Chalakova Z., Rayment J., Ellis B.C.S., Stender E.G.P., van Vugt J.J.F.A., et al. (2024) Project MinE ALS Sequencing Consortium TDP-43 is a Master Regulator of Paraspeckle Condensation. SSRN https://ssrn.com/abstract=4721338or10.2139/ssrn.4721338

Huang, W.P., Ellis, B.C.S., Hodgson, R.E., Sanchez Avila, A., Kumar, V., Rayment, J., Moll, T., and Shelkovnikova, T.A. (2024). Stress-induced TDP-43 nuclear condensation causes splicing loss of function and STMN2 depletion. Cell Rep 43, 114421.

Huang, Y., Su, R., Sheng, Y., Dong, L., Dong, Z., Xu, H., Ni, T., Zhang, Z.S., Zhang, T., Li, C., et al. (2019). Small-Molecule Targeting of Oncogenic FTO Demethylase in Acute Myeloid Leukemia. Cancer Cell 35, 677–691 e610.

Humphrey, J., Birsa, N., Milioto, C., McLaughlin, M., Ule, A.M., Robaldo, D., Eberle, A.B., Krauchi, R., Bentham, M., Brown, A.L., et al. (2020). FUS ALS-causative mutations impair FUS autoregulation and splicing factor networks through intron retention. Nucleic Acids Res 48, 6889–6905.

Jain, S., Wheeler, J.R., Walters, R.W., Agrawal, A., Barsic, A., and Parker, R. (2016). ATPase-Modulated Stress Granules Contain a Diverse Proteome and Substructure. Cell 164, 487–498.

Khoroshkin, M., Buyan, A., Dodel, M., Navickas, A., Yu, J., Trejo, F., Doty, A., Baratam, R., Zhou, S., Lee, S.B., et al. (2024). Systematic identification of post-transcriptional regulatory modules. Nat Commun 15, 7872.

Koike, Y., Pickles, S., Estades Ayuso, V., Jansen-West, K., Qi, Y.A., Li, Z., Daughrity, L.M., Yue, M., Zhang, Y.J., Cook, C.N., et al. (2023). TDP-43 and other hnRNPs regulate cryptic exon inclusion of a key ALS/FTD risk gene, UNC13A. PLoS Biol 21, e3002028.

Konigs, V., de Oliveira Freitas Machado, C., Arnold, B., Blumel, N., Solovyeva, A., Lobbert, S., Schafranek, M., Ruiz De Los Mozos, I., Wittig, I., McNicoll, F., et al. (2020). SRSF7 maintains its homeostasis through the expression of Split-ORFs and nuclear body assembly. Nat Struct Mol Biol 27, 260–273.

Korobeynikov, V.A., Lyashchenko, A.K., Blanco-Redondo, B., Jafar-Nejad, P., and Shneider, N.A. (2022). Antisense oligonucleotide silencing of FUS expression as a therapeutic approach in amyotrophic lateral sclerosis. Nat Med 28, 104–116.

Kwiatkowski, T.J., Jr., Bosco, D.A., Leclerc, A.L., Tamrazian, E., Vanderburg, C.R., Russ, C., Davis, A., Gilchrist, J., Kasarskis, E.J., Munsat, T., et al. (2009). Mutations in the FUS/TLS gene on chromosome 16 cause familial amyotrophic lateral sclerosis. Science 323, 1205–1208.

Lashley, T., Rohrer, J.D., Bandopadhyay, R., Fry, C., Ahmed, Z., Isaacs, A.M., Brelstaff, J.H., Borroni, B., Warren, J.D., Troakes, C., et al. (2011). A comparative clinical, pathological, biochemical and genetic study of fused in sarcoma proteinopathies. Brain 134, 2548–2564.

Lee, E.S., Smith, H.W., Wang, Y.E., Ihn, S.S., Scalize de Oliveira, L., Kejiou, N.S., Liang, Y.L., Nabeel-Shah, S., Jomphe, R.Y., Pu, S., et al. (2025). N-6-methyladenosine (m6A) promotes the nuclear retention of mRNAs with intact 5’ splice site motifs. Life Sci Alliance 8.

Lee, J.H., Wang, R., Xiong, F., Krakowiak, J., Liao, Z., Nguyen, P.T., Moroz-Omori, E.V., Shao, J., Zhu, X., Bolt, M.J., et al. (2021). Enhancer RNA m6A methylation facilitates transcriptional condensate formation and gene activation. Molecular cell 81, 3368–3385 e3369.

Liang, Z., Ye, H., Ma, J., Wei, Z., Wang, Y., Zhang, Y., Huang, D., Song, B., Meng, J., Rigden, D.J., et al. (2024). m6A-Atlas v2.0: updated resources for unraveling the N6-methyladenosine (m6A) epitranscriptome among multiple species. Nucleic Acids Res 52, D194–D202.

Lin, C.L., Leu, S., Lu, M.C., and Ouyang, P. (2004). Over-expression of SR-cyclophilin, an interaction partner of nuclear pinin, releases SR family splicing factors from nuclear speckles. Biochemical and biophysical research communications 321, 638–647.

Ling, S.C., Dastidar, S.G., Tokunaga, S., Ho, W.Y., Lim, K., Ilieva, H., Parone, P.A., Tyan, S.H., Tse, T.M., Chang, J.C., et al. (2019). Overriding FUS autoregulation in mice triggers gain-of-toxic dysfunctions in RNA metabolism and autophagy-lysosome axis. Elife 8.

Lopez-Erauskin, J., Tadokoro, T., Baughn, M.W., Myers, B., McAlonis-Downes, M., Chillon-Marinas, C., Asiaban, J.N., Artates, J., Bui, A.T., Vetto, A.P., et al. (2018). ALS/FTD-Linked Mutation in FUS Suppresses Intra-axonal Protein Synthesis and Drives Disease Without Nuclear Loss-of-Function of FUS. Neuron 100, 816–830.

Lorenz, R., Bernhart, S.H., Honer Zu Siederdissen, C., Tafer, H., Flamm, C., Stadler, P.F., and Hofacker, I.L. (2011). ViennaRNA Package 2.0. Algorithms Mol Biol 6, 26.

Ma, W., Zhen, G., Xie, W., and Mayr, C. (2021). In vivo reconstitution finds multivalent RNA-RNA interactions as drivers of mesh-like condensates. Elife 10.

Martinez, F.J., Pratt, G.A., Van Nostrand, E.L., Batra, R., Huelga, S.C., Kapeli, K., Freese, P., Chun, S.J., Ling, K., Gelboin-Burkhart, C., et al. (2016). Protein-RNA Networks Regulated by Normal and ALS-Associated Mutant HNRNPA2B1 in the Nervous System. Neuron 92, 780–795.

Mathews, D.H., Disney, M.D., Childs, J.L., Schroeder, S.J., Zuker, M., and Turner, D.H. (2004). Incorporating chemical modification constraints into a dynamic programming algorithm for prediction of RNA secondary structure. Proc Natl Acad Sci U S A 101, 7287–7292.

Mauger, O., Lemoine, F., and Scheiffele, P. (2016). Targeted Intron Retention and Excision for Rapid Gene Regulation in Response to Neuronal Activity. Neuron 92, 1266–1278.

Mazille, M., Buczak, K., Scheiffele, P., and Mauger, O. (2022). Stimulus-specific remodeling of the neuronal transcriptome through nuclear intron-retaining transcripts. EMBO J 41, e110192.

Mellacheruvu, D., Wright, Z., Couzens, A.L., Lambert, J.P., St-Denis, N.A., Li, T., Miteva, Y.V., Hauri, S., Sardiu, M.E., Low, T.Y., et al. (2013). The CRAPome: a contaminant repository for affinity purification-mass spectrometry data. Nat Methods 10, 730–736.

Mitchell, J.C., McGoldrick, P., Vance, C., Hortobagyi, T., Sreedharan, J., Rogelj, B., Tudor, E.L., Smith, B.N., Klasen, C., Miller, C.C., et al. (2013). Overexpression of human wild-type FUS causes progressive motor neuron degeneration in an age- and dose-dependent fashion. Acta neuropathologica 125, 273–288.

Moens, T.G., Da Cruz, S., Neumann, M., Shelkovnikova, T.A., Shneider, N.A., and Van Den Bosch, L. (2025). Amyotrophic lateral sclerosis caused by FUS mutations: advances with broad implications. Lancet Neurol 24, 166–178.

Nolan, M., Talbot, K., and Ansorge, O. (2016). Pathogenesis of FUS-associated ALS and FTD: insights from rodent models. Acta Neuropathol Commun 4, 99.

Patel, A., Mitrea, D., Namasivayam, V., Murcko, M.A., Wagner, M., and Klein, I.A. (2022). Principles and functions of condensate modifying drugs. Front Mol Biosci 9, 1007744.

Patil, D.P., Chen, C.K., Pickering, B.F., Chow, A., Jackson, C., Guttman, M., and Jaffrey, S.R. (2016). m(6)A RNA methylation promotes XIST-mediated transcriptional repression. Nature 537, 369–373.

Ratti, A., and Buratti, E. (2016). Physiological Functions and Pathobiology of TDP-43 and FUS/TLS proteins. J Neurochem 138 Suppl 1:95–111.

Ries, R.J., Zaccara, S., Klein, P., Olarerin-George, A., Namkoong, S., Pickering, B.F., Patil, D.P., Kwak, H., Lee, J.H., and Jaffrey, S.R. (2019). m(6)A enhances the phase separation potential of mRNA. Nature 571, 424–428.

Rogelj, B., Easton, L.E., Bogu, G.K., Stanton, L.W., Rot, G., Curk, T., Zupan, B., Sugimoto, Y., Modic, M., Haberman, N., et al. (2012). Widespread binding of FUS along nascent RNA regulates alternative splicing in the brain. Sci Rep 2, 603.

Sabatelli, M., Moncada, A., Conte, A., Lattante, S., Marangi, G., Luigetti, M., Lucchini, M., Mirabella, M., Romano, A., Del Grande, A., et al. (2013). Mutations in the 3’ untranslated region of FUS causing FUS overexpression are associated with amyotrophic lateral sclerosis. Human molecular genetics 22, 4748–4755.

Sanjuan-Ruiz, I., Govea-Perez, N., McAlonis-Downes, M., Dieterle, S., Megat, S., Dirrig-Grosch, S., Picchiarelli, G., Piol, D., Zhu, Q., Myers, B., et al. (2021). Wild-type FUS corrects ALS-like disease induced by cytoplasmic mutant FUS through autoregulation. Mol Neurodegener 16, 61.

Scekic-Zahirovic, J., Sendscheid, O., El Oussini, H., Jambeau, M., Sun, Y., Mersmann, S., Wagner, M., Dieterle, S., Sinniger, J., Dirrig-Grosch, S., et al. (2016). Toxic gain of function from mutant FUS protein is crucial to trigger cell autonomous motor neuron loss. EMBO J 35, 1077–1097.

Sharma, A., Lyashchenko, A.K., Lu, L., Nasrabady, S.E., Elmaleh, M., Mendelsohn, M., Nemes, A., Tapia, J.C., Mentis, G.Z., and Shneider, N.A. (2016). ALS-associated mutant FUS induces selective motor neuron degeneration through toxic gain of function. Nat Commun 7, 10465.

Shelkovnikova, T.A., An, H., Skelt, L., Tregoning, J.S., Humphreys, I.R., and Buchman, V.L. (2019). Antiviral Immune Response as a Trigger of FUS Proteinopathy in Amyotrophic Lateral Sclerosis. Cell Rep 29, 4496–4508 e4494.

Shelkovnikova, T.A., Dimasi, P., Kukharsky, M.S., An, H., Quintiero, A., Schirmer, C., Buee, L., Galas, M.C., and Buchman, V.L. (2017). Chronically stressed or stress-preconditioned neurons fail to maintain stress granule assembly. Cell Death Dis 8, e2788.

Shelkovnikova, T.A., Kukharsky, M.S., An, H.Y., Dimasi, P., Alexeeva, S., Shabir, O., Heath, P.R., and Buchman, V.L. (2018). Protective paraspeckle hyper-assembly downstream of TDP-43 loss of function in amyotrophic lateral sclerosis. Mol Neurodegener 13.

Shelkovnikova, T.A., Robinson, H.K., Southcombe, J.A., Ninkina, N., and Buchman, V.L. (2014a). Multistep process of FUS aggregation in the cell cytoplasm involves RNA-dependent and RNA-independent mechanisms. Human molecular genetics 23, 5211–5226.

Shelkovnikova, T.A., Robinson, H.K., Troakes, C., Ninkina, N., and Buchman, V.L. (2014b). Compromised paraspeckle formation as a pathogenic factor in FUSopathies. Human molecular genetics 23, 2298–2312.

Sung, H.M., Schott, J., Boss, P., Lehmann, J.A., Hardt, M.R., Lindner, D., Messens, J., Bogeski, I., Ohler, U., and Stoecklin, G. (2023). Stress-induced nuclear speckle reorganization is linked to activation of immediate early gene splicing. J Cell Biol 222.

Suzuki, K., Bose, P., Leong-Quong, R.Y., Fujita, D.J., and Riabowol, K. (2010). REAP: A two minute cell fractionation method. BMC Res Notes 3, 294.

Takakuwa, H., Yamazaki, T., Souquere, S., Adachi, S., Yoshino, H., Fujiwara, N., Yamamoto, T., Natsume, T., Nakagawa, S., Pierron, G., et al. (2023). Shell protein composition specified by the lncRNA NEAT1 domains dictates the formation of paraspeckles as distinct membraneless organelles. Nat Cell Biol 25, 1664–1675.

Vance, C., Rogelj, B., Hortobagyi, T., De Vos, K.J., Nishimura, A.L., Sreedharan, J., Hu, X., Smith, B., Ruddy, D., Wright, P., et al. (2009). Mutations in FUS, an RNA processing protein, cause familial amyotrophic lateral sclerosis type 6. Science 323, 1208–1211.

Wang, X., Liu, C., Zhang, S., Yan, H., Zhang, L., Jiang, A., Liu, Y., Feng, Y., Li, D., Guo, Y., et al. (2021). N(6)-methyladenosine modification of MALAT1 promotes metastasis via reshaping nuclear speckles. Dev Cell 56, 702–715 e708.

Widagdo, J., Anggono, V., and Wong, J.J. (2022). The multifaceted effects of YTHDC1-mediated nuclear m(6)A recognition. Trends Genet 38, 325–332.

Wu, J., Xiao, Y., Liu, Y., Wen, L., Jin, C., Liu, S., Paul, S., He, C., Regev, O., and Fei, J. (2024). Dynamics of RNA localization to nuclear speckles are connected to splicing efficiency. Sci Adv 10, eadp7727.

Yankova, E., Blackaby, W., Albertella, M., Rak, J., De Braekeleer, E., Tsagkogeorga, G., Pilka, E.S., Aspris, D., Leggate, D., Hendrick, A.G., et al. (2021). Small-molecule inhibition of METTL3 as a strategy against myeloid leukaemia. Nature 593, 597–601.

Yap, K., Chung, T.H., and Makeyev, E.V. (2022a). Analysis of RNA-containing compartments by hybridization and proximity labeling in cultured human cells. STAR Protoc 3, 101139.

Yap, K., Chung, T.H., and Makeyev, E.V. (2022b). Hybridization-proximity labeling reveals spatially ordered interactions of nuclear RNA compartments. Molecular cell 82, 463–478 e411.

Yeo, G., and Burge, C.B. (2004). Maximum entropy modeling of short sequence motifs with applications to RNA splicing signals. J Comput Biol 11, 377–394.

Zhou, Y., Liu, S., Liu, G., Ozturk, A., and Hicks, G.G. (2013). ALS-associated FUS mutations result in compromised FUS alternative splicing and autoregulation. PLoS Genet 9, e1003895.

Zhou, Y., Zeng, P., Li, Y.H., Zhang, Z., and Cui, Q. (2016). SRAMP: prediction of mammalian N6-methyladenosine (m6A) sites based on sequence-derived features. Nucleic Acids Res 44, e91.

Ziff, O.J., Harley, J., Wang, Y., Neeves, J., Tyzack, G., Ibrahim, F., Skehel, M., Chakrabarti, A.M., Kelly, G., and Patani, R. (2023). Nucleocytoplasmic mRNA redistribution accompanies RNA binding protein mislocalization in ALS motor neurons and is restored by VCP ATPase inhibition. Neuron 111, 3011–3027 e3017.

