## Supplementary material for "FUS post-transcriptional splicing is autoregulated via RNA condensation with therapeutic potential for ALS-FUS"

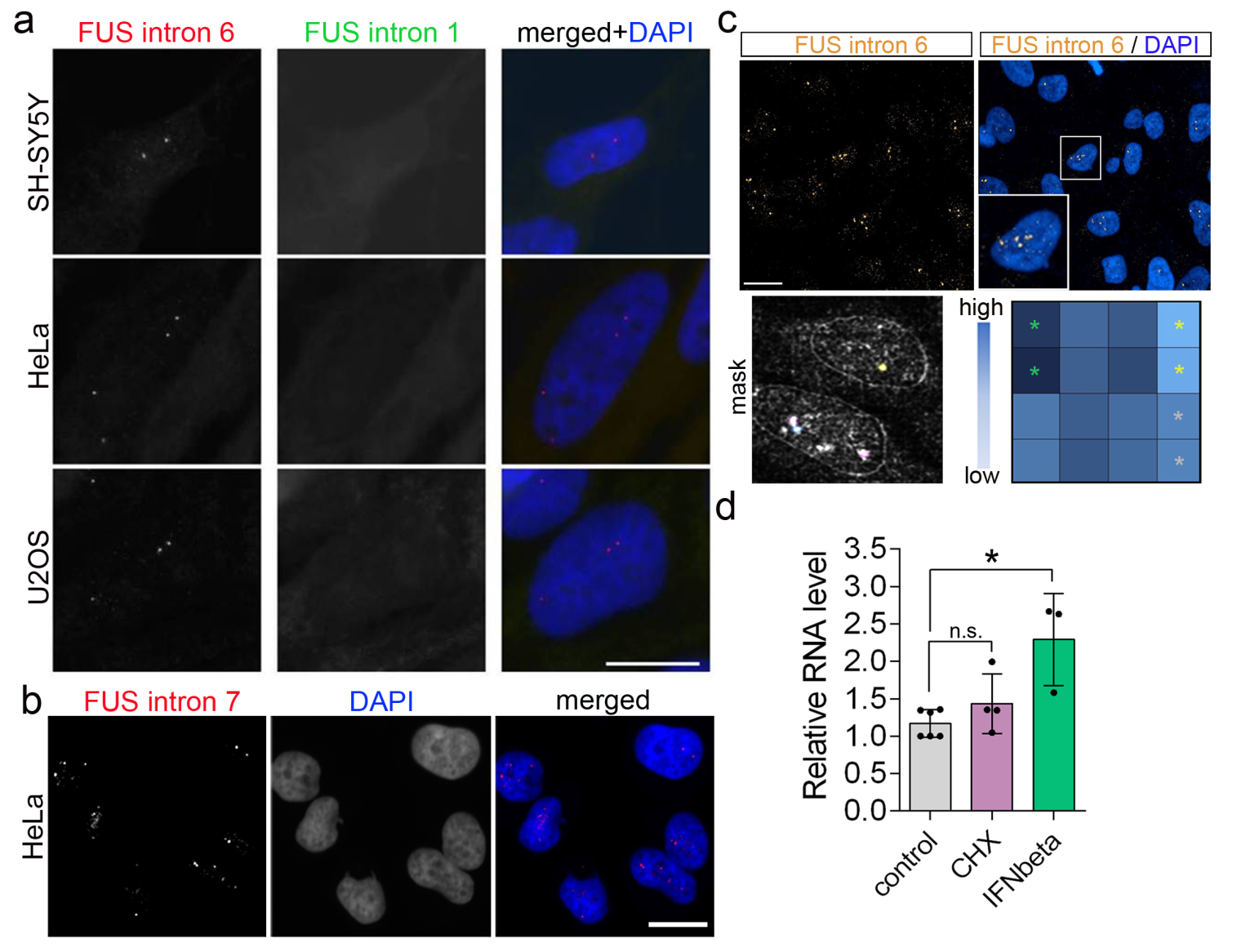

**Supplementary Figure 1. FUSint6&7-RNA foci characterisation.**

**a** FUSint6&7-RNA foci are present in different human cell lines and are not detected by a probe mapping to FUS intron 1. Scale bar, 10 µm.

**b** FUSint6&7-RNA foci detection using FUS intron 7-specific probe. Scale bar, 15 µm.

**c** Automated quantification assay for FUSint6&7-RNA foci analysis. Opera Phenix HCS confocal system and a custom quantification pipeline on Harmony were used. Yellow, grey and green dots in the heatmap (foci number) indicate no probe control, DMSO control, and IFNbeta (positive control), respectively.

**d** FUSint6&7-RNA is insensitive to NMD and is upregulated by IFNbeta treatment, as demonstrated by qRT-PCR analysis with FUS intron 6-specific primers. N=4-5, *p<0.05, Kruskal-Wallis with Dunn’s test.

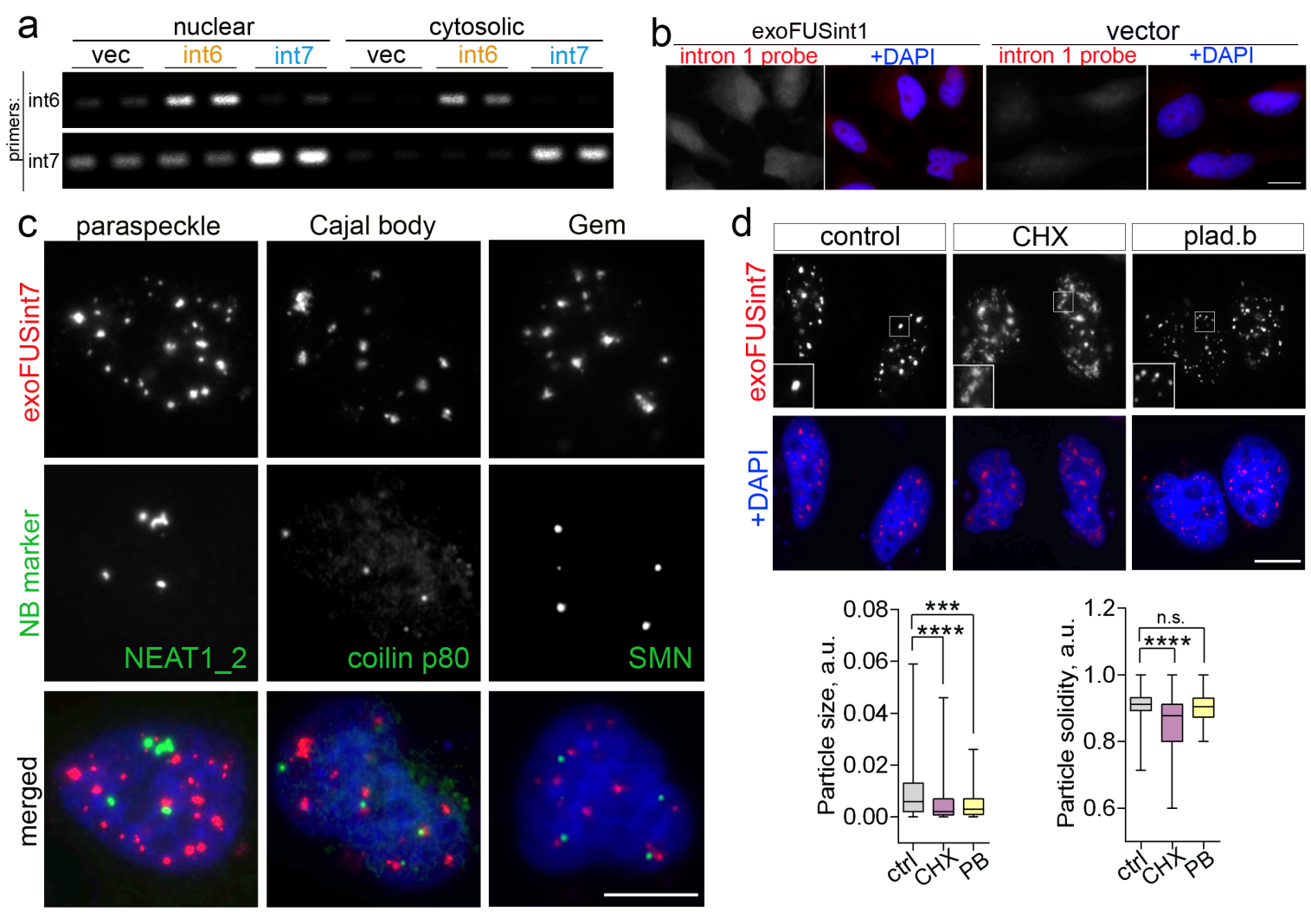

**Supplementary Figure 2. Characterisation of ectopically expressed FUS introns.**

**a** Ectopically expressed exoFUSint6 and -7 are enriched in the nuclear fraction. RT-PCR analysis was performed on RNA isolated from nuclear and cytosolic fractions of cells expressing vector control or respective introns.

**b** FUS intron 1 displays diffuse distribution when overexpressed. Scale bar, 10 μm.

**c** ExoFUSint7 condensates do not overlap with known nuclear bodies. Representative images are shown. Scale bar, 10 μm.

**d** ExoFUSint7 condensates are sensitive to changes in RNA metabolism/processing, similar to the endogenous FUSint6&7-RNA condensates. Cells were treated with pladienolide B or CHX for 4 h. Note that CHX leads to dissipation of condensates (decreased individual particle size and solidity). Representative images and quantification are shown. ~100 nuclear particles (>100px in size) were analysed per condition. ***p<0.001, ****p<0.0001, Kruskal-Wallis with Dunn’s test. Scale bar, 5 μm.

HeLa cells were used for these studies.

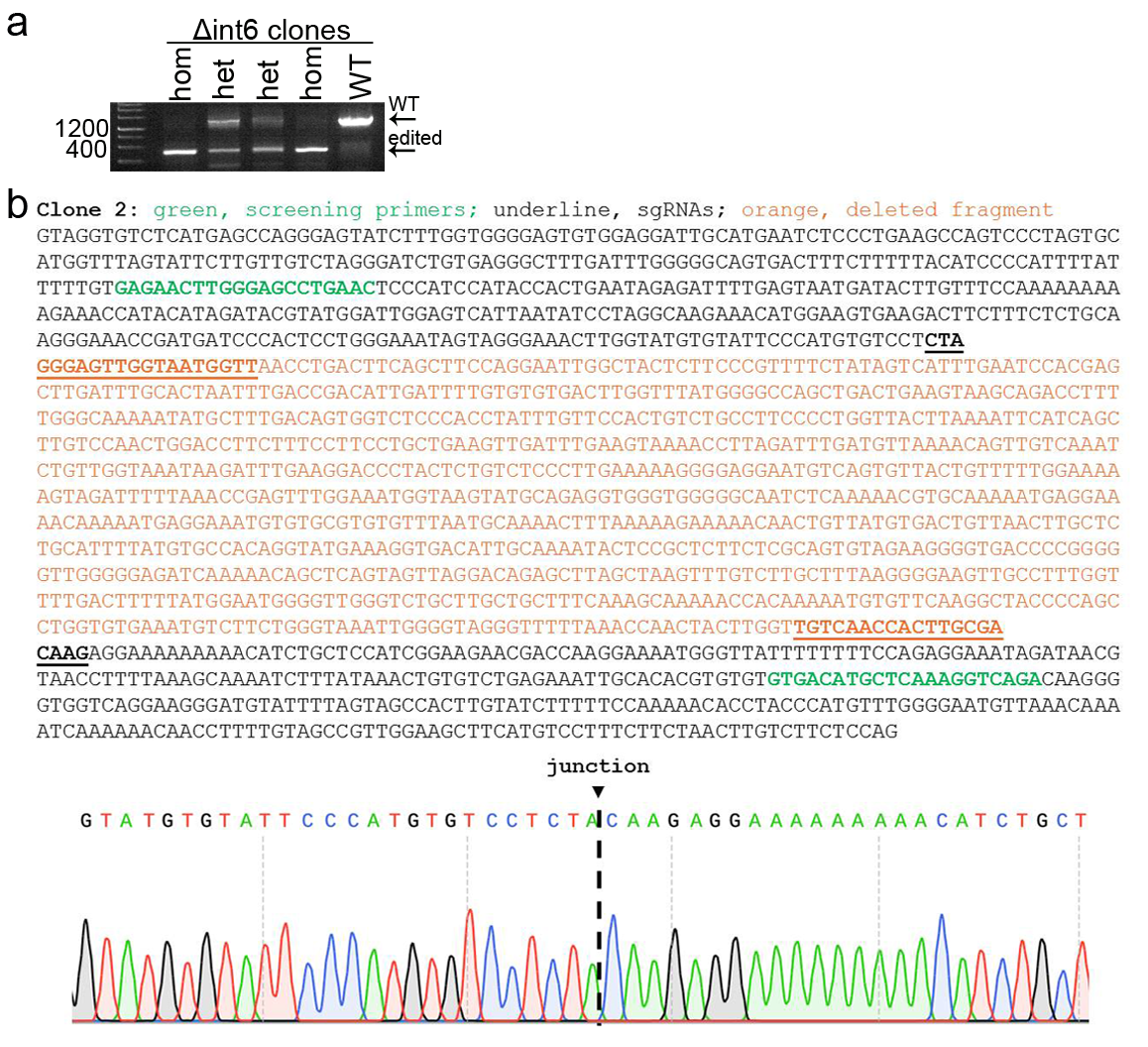

**Supplementary Figure 3. Characterisation of cell clones with FUS intron 6 deletion.**

**a** Confirmation of homo- and heterozygous Δint6 clones by PCR on genomic DNA.

**b** Confirmation of the expected editing outcome by sequencing in a homozygous clone. The edited junction was sequenced using the PCR product corresponding to the edited band as in *a*.

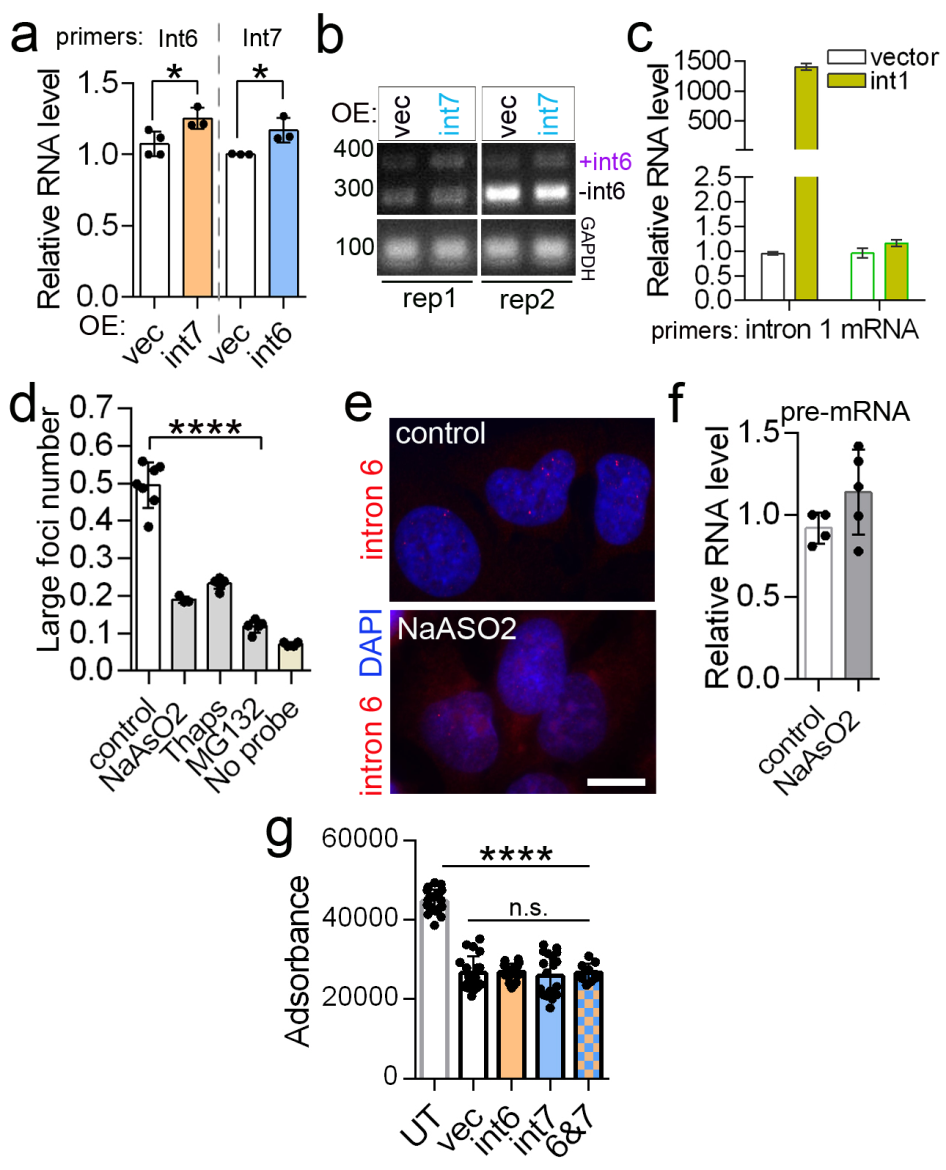

**Supplementary Figure 4. Characterisation of FUS intron effects on FUS RNA processing and the effect of stress.**

**a,b** Ectopic expression of exoFUSint6- and -7 upregulates endogenous FUSint6&7-RNA. RNA expression was measured by qRT-PCR (a) or RT-PCR (b). N=3-4, *p<0.05, Mann-Whitney *U* test.

**c** Ectopic expression of FUS intron 1 (exoFUSint1) does not affect FUS mRNA levels. Analysis was done by qRT-PCR. N=3.

**d,e** FUSint6&7-RNA condensates are reduced in response to cellular stress. Cells were treated with MG132 and thapsigargin for 4 h, or treated with NaAsO2 for 1 h and left to recover for 3 h (total stress duration = 4 h) and analysed by automated imaging (d). N=3-6 (individual wells), ****p<0.0001, one-way ANOVA with Dunnett’s test. Loss of condensates was also confirmed by high-resolution imaging (e). Scale bar, 10 μm.

**f** FUS transcription is not affected by oxidative stress. Levels of FUS pre-mRNA were analysed using qRT-PCR with intron 1-specific primers during the recovery from NaAsO2 (1 h stress+3 h recovery). N=4-5.

**g** Ectopic expression of FUS introns 6/7 is not cytotoxic. Survival of cells transfected with a vector control or exoFUSint6 or -7 expression constructs was analysed using a resazuin-based assay. 20 wells from 2 independent experiments were used for analysis. ****p<0.0001, one-way ANOVA with Dunnett’s post-hoc test; UT, untransfected; n.s., non-significant.

HeLa cells were used in these experiments.

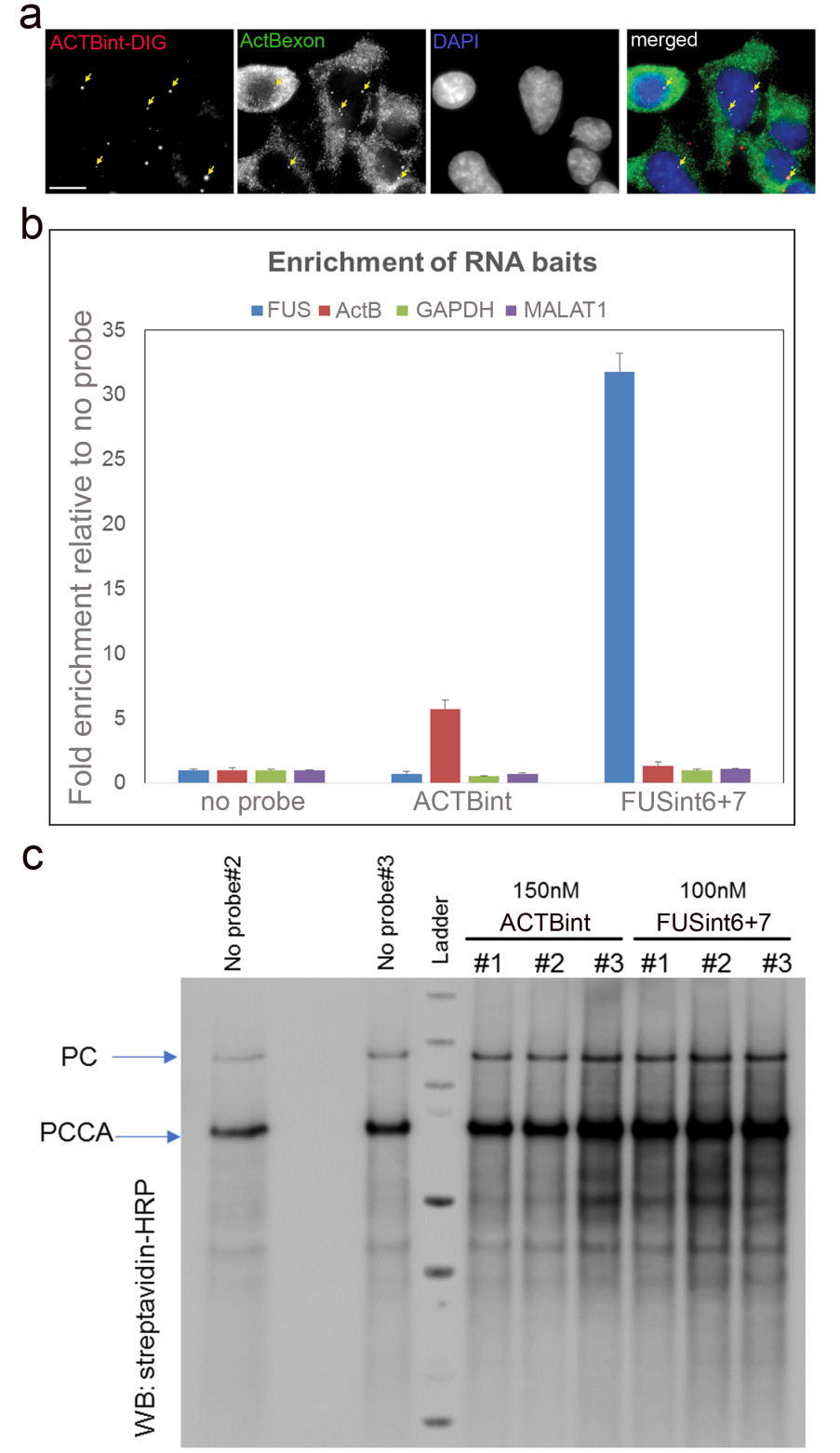

**Supplementary Figure 5. Quality control for HyPro-MS.**

**a** HyPro-FISH with ACTB probes. Scale bar, 10 µm.

**b** RNA bait enrichment analysis of the experimental triplicates after HyPro labelling and pulldown. qRT-PCR primers are given in Supplementary Table S4.

**c** Biotinylation analysis of the experimental triplicates for FUSint6&7-RNA, ACTB introns and no-probe conditions by western blot. Arrows indicate PC and PCCA – endogenous proteins that are frequently biotinylated.

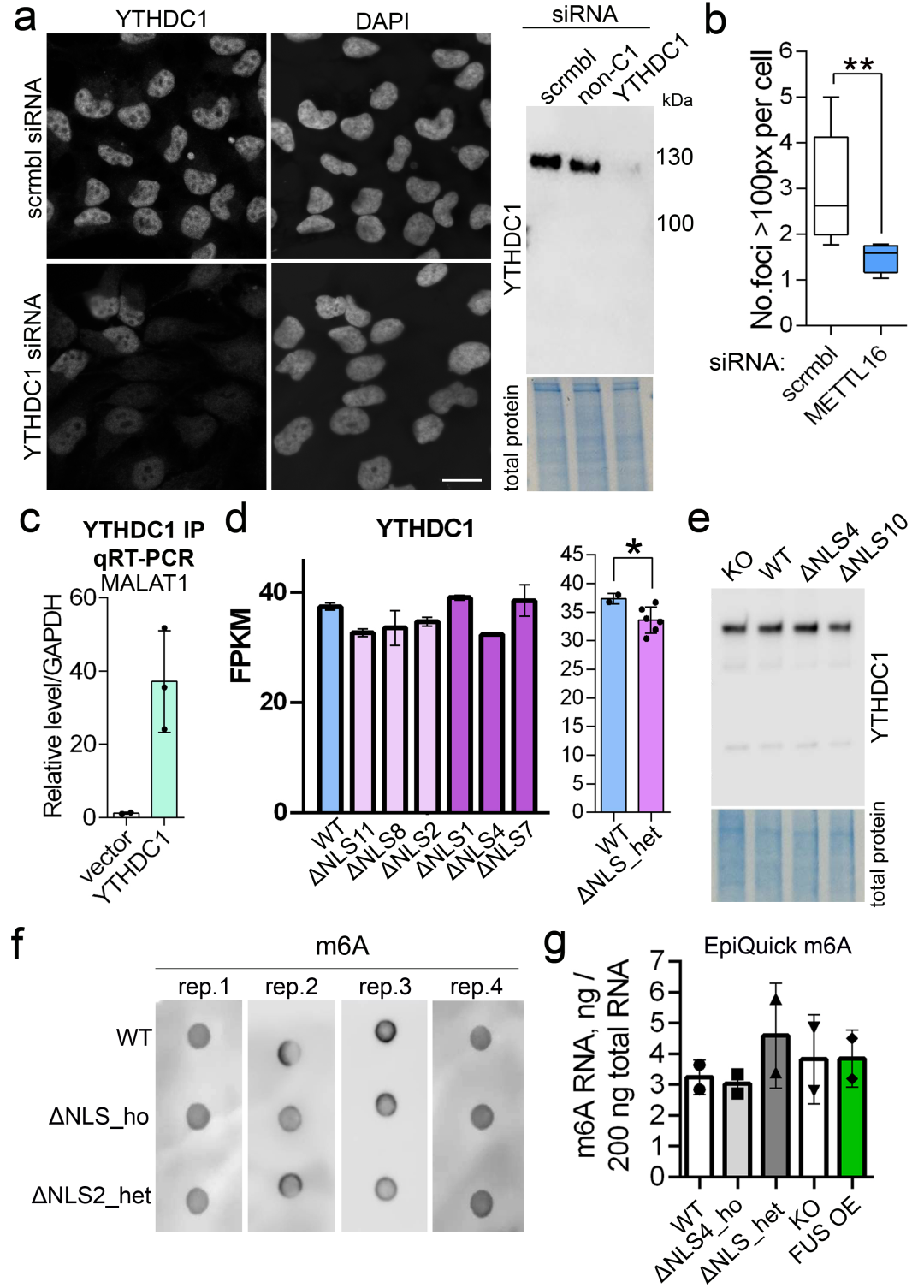

**Supplementary Figure 6. m6A/YTHDC1 in FUSint6&7-RNA regulation in WT and mutant FUS expressing cells.**

**a** Confirmation of YTHDC1 siRNA-mediated knockdown by immunostaining and western blot in HeLa cells. Scale bar, 20 μm.

**b** METTL16 depletion reduces FUSint6&7-RNA condensate assembly. Quantification was done using FUS intron 6 RNA-FISH in HeLa cells. 155 and 102 cells (4-5 FoV) were analysed for scrambled and METTL16 siRNAs, respectively, from a representative experiment. **p<0.01, Mann-Whitney *U* test.

**c** Efficient MALAT1 pulldown in a RIP experiment with YTHDC1-Flag and Flag-Trap beads. N=3.

**d** YTHDC1 mRNA downregulation in FUSΔNLS lines. RNAseq data were from An et al., 2019. Right graph shows combined data for the 3 heterozygous lines. *p<0.05, Mann-Whitney *U* test.

**e** YHTDC1 protein levels are not affected in FUSΔNLS lines. Representative western blot is shown.

**f,g** Total m6A levels are not altered in FUSΔNLS lines. Dot blot with an m6A antibody (f) and EpiQuick ELISA-type assay (g) were used. N=2-4.

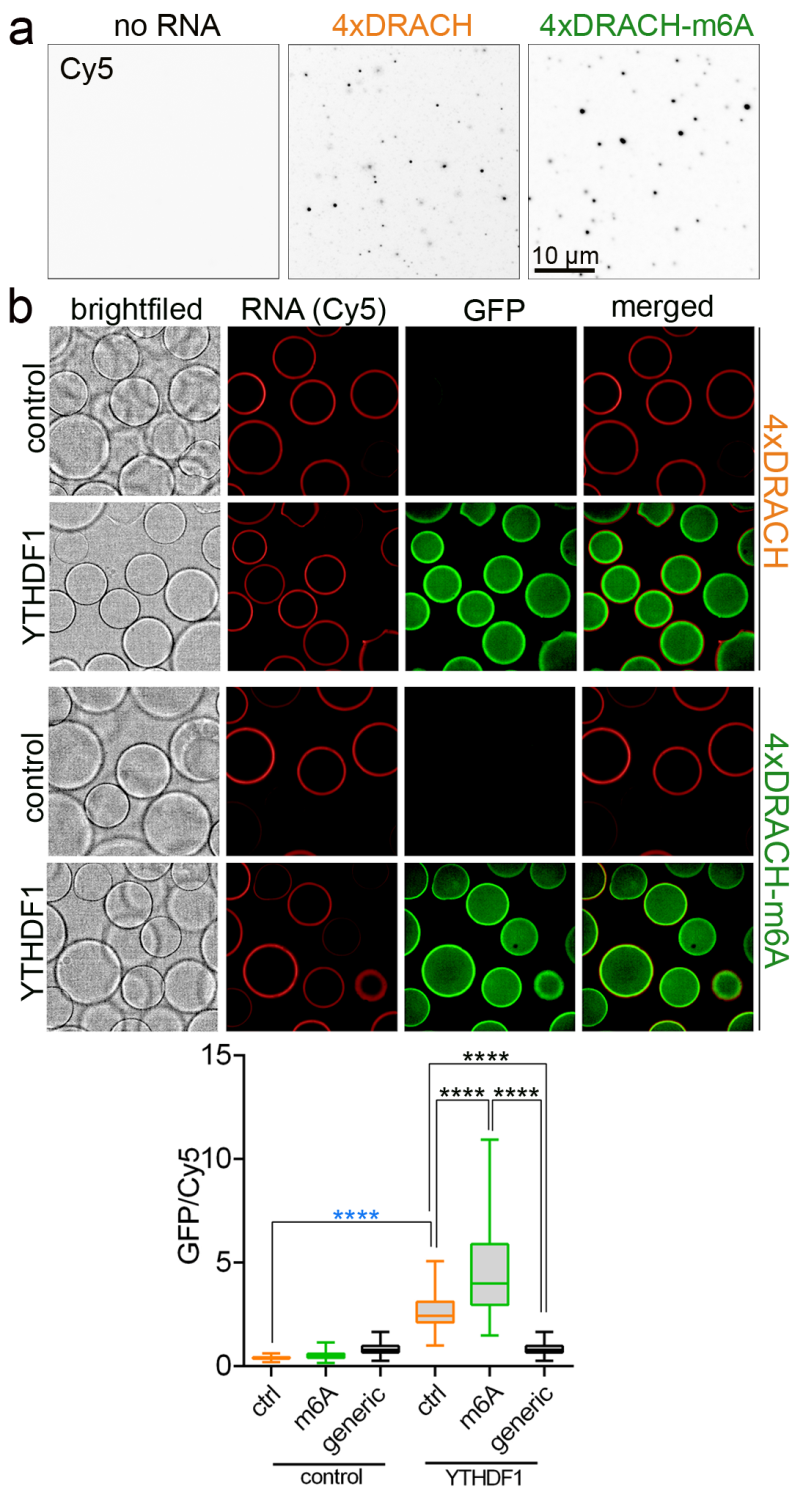

**Supplementary Figure 7. m6A effect on RNA condensation and a positive control for CONA.**

**a** m6A promotes RNA condensation. Condensate formation by 4xDRACH and 4xDRACH RNA oligonucleotides (Cy5-labelled), in the presence of recombinant FUS protein. Condensates were imaged using Cy5 fluorescence after sedimentation and fixation on cover glasses.

**b** Validation of CONA use for methylated RNA studies using an m6A reader, YTHDF1. Lysates of cells expressing either GFP or YTHDF1-GFP were used. Generic RNA corresponds to a mix of three RNA oligonucleotides with diverse sequences: (UG)_15_, (AUG)_12_, Clip34nt, mixed at an equal ratio. Representative images and quantification are shown. 80-100 beads were analysed per condition. ****p<0.0001, Mann-Whitney *U* test (ctrl RNA with GFP vs. ctrl RNA with YTHDF1; blue asterisks) or Kruskal-Wallis with Dunn’s test (YTHDF1: ctrl RNA vs. m6A RNA and generic RNA; black asterisks).

**Supplementary Table 1. MaxEntScan analysis of FUS splice sites.**

| Exon | Acc Seq | Don Seq | Acc MaxEnt | Don MaxEnt | Acc WMM | Don WMM |
| --- | --- | --- | --- | --- | --- | --- |
| Exon 1 | NA | ACGgtaggt | NA | 10.15 | NA | 7.99 |
| Exon 2 | attgtatttttcttttgcagATT | AAGgtgagt | 11.61 | 10.47 | 13.39 | 11.33 |
| Exon 3 | ctggttttccttttatttagCTA | ACAgtgagt | 7.26 | 8.34 | 10.51 | 5.81 |
| Exon 4 | cctttttcttatcctggtagCAG | AAGgtacgg | 6.31 | 10.26 | 10.03 | 8.79 |
| Exon 5 | tttttgtttgttttccctagTTA | GAGgtgaga | 9.67 | 7.66 | 13.05 | 9.14 |
| Exon 6 | cattctttcttttctcacagGTA | GGGgtaggt | 12.23 | 6.59 | 15.77 | 7.22 |
| Exon 7 | ttctaacttgtcttctccagCGG | GTGgtaagt | 8.22 | 10.36 | 10.4 | 9.83 |
| Exon 8 | ttttttccatgtcactaaagGCC | CCGgtgagt | 5.03 | 10.9 | 7.08 | 9.35 |
| Exon 9 | tttctctgttcaacaagcagAAC | AAGgtactt | 4.9 | 8.4 | 4.36 | 5.82 |
| Exon 10 | cctcattttgctttcttcagACA | ATGgtatgt | 7.91 | 8.35 | 11.97 | 7.72 |
| Exon 11 | atgattttttgtttctctagGTA | GAGgtgagg | 11.14 | 8.41 | 13.15 | 9.56 |
| Exon 12 | acttggtctatctgcattagGAC | TCCgtgagt | 5.81 | 8.93 | 5.45 | 4.46 |
| Exon 13 | ttgtcttcctttctccttagCAC | TGGgtaaga | 9.74 | 8.91 | 13.66 | 7.41 |
| Exon 14 | tcttgtttcttttgtcctagGGG | CAGgtaaga | 9.76 | 10.77 | 13.37 | 11.73 |
| Exon 15 | ttttttttttttttttgcagGGG | NA | 11.95 | NA | 18.36 | NA |

**retained region flanking sites*

**Supplementary Table 2. HyPro-MS analysis of endogenous FUSint6&7-RNA condensates.**

*Available as Excel file.*

**Supplementary Table 3. Analysis of m6A modifications on FUS RNA.**

*Available as Excel file.*

**Supplementary Table 4. Primers used in the study.**

| **qPCR and PCR primers – main** | | |
| --- | --- | --- |
| **Primer** | **Forward** | **Reverse** |
| FUS_ex5_int6 (PCR) | 5'-GCTATGGACAGCAGCAAAGC-3’ | 5'-CAGTCAGCTGGCCCCATAAA-3’ |
| FUS_int7_ex11 (PCR) | 5’-GTTTCGGGGAAACAACACGG-  3’ | 5’-CCGGAGAATTCTTTACCATCAAACC  -3’ |
| FUS_int6 (qPCR) | 5'-TTTATGGGGCCAGCTGACTG-3’ | 5'-GTAACCAGGGGAAGGCAGAC-3’ |
| FUS_int7 (qPCR) | 5'-TGTGTGCTAACCTGGAGCAG-3’ | 5'-TCCAAGGCCTACTAGACCCC-3’ |
| FUS_total (int6/7+mRNA) (qPCR) | 5’-GGAACTCAGTCAACTCCCCA-3’ | 5’-TACCGTAACTTCCCGAGGTG-3’ |
| FUS_int 1/pre-mRNA (qPCR) | 5’-AAGCCGCGGAGAAGAGTAA-3’ | 5’-AAGAAAAGACTTCCCGCCCC-3’ |
| FUS mRNA only (qPCR) | 5’-CGGCGGTGGTGGTTACAA-3’ | 5’-GTCCCGAGGGCCACCAAAT-3’ |
| FUS_ex6_for (PCR) | 5'-TCCTCCATGAGTAGTGGTGGT-3’ | *na* |
| FUS_int6_rev (PCR) | *na* | 5'-GTTCAGGCTCCCAAGTTCTC-3’ |
| FUS_ex8/9_rev (PCR) | *na* | 5'-GTCTGAATTATCCTGTTCGGAGTC-3’ |
| FUSint6 (cloning) | 5’-gatctcgagGTGGCATGGGgtaggtgtct-3’ | 5’-cgcggtaccTCACTTCCGctggagaagac-3’ |
| FUSint7 (cloning) | 5’-gatctcgagCAATAAATTTGGTGgtaagtgaacagagtttc-3’ | 5’-cgcggtaccCCCGAGGGCctttagtgaca-3’ |
| FUSint1 (cloning) | 5’-gcagtcgacGCCTCAAACGgtaggtaagg-3 | 5’-ggtggatccCTTGTTGGGTATAATctgcaaaaga-3’ |
| Δint6 cell line (screening by PCR) | 5’-GAGAACTTGGGAGCCTGAAC-3’ | 5’-TCTGACCTTTGAGCATGTCAC-3’ |
| **qPCR primers – HyPro QC** | | |
| hFUS | 5’-GGTGGTCAGGAAGGGATGTA-3’ | 5’-ACCACCAAATTTATTGAAGCCAC-3’ |
| hACTB | 5’-TGGCACCACACCTTCTACAA-3’ | 5’-AACGGCAGAAGAGAGAACCA-3’ |
| hGAPDH | 5’-CCTGACCTGCCGTCTAGAAA-3’ | 5’-CCCTGTTGCTGTAGCCAAAT-3’ |
| hMALAT1 | 5’-TGATGGCCTAGATGCAGAGAA-3’ | 5’-GAGATGGACATTGCCTCTTCA-3’ |

**Supplementary Table 5. RNA *in situ* hybridisation probes used in the study.**

*Available as Excel file.*
